# Anode surface bioaugmentation enhances deterministic biofilm assembly in microbial fuel cells

**DOI:** 10.1101/2020.02.17.951574

**Authors:** Keren Yanuka-Golub, Vadim Dubinsky, Elisa Korenblum, Leah Reshef, Maya Ofek-Lalzar, Judith Rishpon, Uri Gophna

## Abstract

Microbial fuel cells (MFCs) are devices that can generate energy while aiding biodegradation of waste through the activity of an electroactive mixed biofilm. Metabolic cooperation is considered essential for MFCs’ efficiency, especially during early-anode colonization. Yet, the specific ecological processes that drive the assembly of an optimized anode-attached community remain unknown. Here, we show, using 16S rRNA gene amplicon and shotgun metagenomic sequencing that bioaugmentation of the anode surface with an electroactive consortium originating from a well-established anodic biofilm, dominated by different *Desulfuromonas* strains, resulted in an extremely rapid voltage generation (reaching maximal voltage within several hours). This was in sharp contrast to the highly stochastic and slower biofilm assembly that occurred when the anode-surface was not augmented. By comparing two inoculation media, wastewater and filtered wastewater, we were able to illustrate two different "source-communities" for newly arriving species that with time colonized the anode surface in a different manner and resulted in dramatically different community assembly processes. Remarkably, an efficient anode colonization process was obtained only if unfiltered wastewater was added, leading to a near-complete replacement of the bioaugmented community by *Geobacter lovleyi*. We propose that anode bioaugmentation reduced stochasticity by creating available niches that were quickly occupied by specific newly-arriving species that positively supported the fast establishment of a highly-functional anode biofilm.

## 1. Introduction

Microbial Fuel Cells (MFC) are electrochemical reactors that exploit microorganisms to promote the generation of electricity through degradation of organic substrates. The operation of MFCs is based on the catalytic activity of an anaerobic electro-active biofilm that develops on the anode’s surface. Bacteria within this biofilm oxidize the organic matter, release electrons extracellularly [via extracellular electron transport (EET)] and use the anode as the terminal electron acceptor (Logan, 2009; Lovley, 2006). EET is a key metabolic processes that facilitates organic matter degradation by anaerobic microbial communities in natural environments under specific redox gradients (Stams et al., 2006). The electroactive biofilm formation process on the anode-surface is critical for an efficient electricity producing MFC (Kumar et al., 2015; Li and Cheng, 2019), and changes in this biofilm consequently strongly affect its electrical output. Hence, the anodic biofilm microbial composition and function has been considered a vital component of the MFC performance, and the metabolic processes that characterize it have been intensively studied (Chae et al. 2009; Beecroft et al. 2012; Commault et al. 2015; Lee et al. 2015; Cui et al. 2016; Hodgson et al. 2016; Paitier et al. 2016; Yanuka-Golub et al., 2016; Ishii et al., 2018).

Establishment of the anodic biofilm, especially in the initial stages of colonization, involves both stochastic and specific selection processes that eventually dictate subsequent biofilm community composition at steady state, defined as a stable maximal current density (Zhang et al., 2019; Zhou et al., 2013). The colonization of MFC anodes is generally regarded as a gradual stochastic process (Beecroft et al., 2012; Chignell et al., 2018; Ishii et al., 2017). Accordingly, our recent results also show that MFCs operated similarly from the same inoculum can substantially vary in lag times (i.e. the time at which current reached a minimal value that is above the baseline value) and startup times (i.e. the time to reach stable maximal current generation) showed different microbial electroactive communities once in steady state. While most studies focus primarily on biofilm formation on clean surfaces (Brislawn et al., 2019; Fillinger et al., 2019; Zhang et al., 2019; Zhou et al., 2013), here we explored the effect of community assembly processes when the anode surface was pre-colonized (bioaugmented) by a defined electroactive consortium (designated as EDC) on MFC startup. Such pre-colonization is expected to enhance deterministic factors relative to stochastic ones in the formation of the mature, anode biofilm at steady state. We show that by augmenting the anode surface with EDC, the microbial performance was dramatically stimulated to produce an outstanding rapid current compared with a slow startup when the anode surface was not applied as such. Importantly, optimal performance was observed by the augmented anode only when supplemented with wastewater, which points to key interspecies-interactions that were dependent on specific newly arriving species provided through the wastewater.

## Materials and Methods

### MFC Setup, experimental design and operation

Single-chamber, air-cathode MFCs were used for the enrichment of electroactive bacteria and suspended-cells (plankton) communities, as previously described (Yanuka-Golub et al., 2016). The experimental setup aimed to directly apply on the anode surface an electroactive-consortium, EDC, composed of >50% *Desulfuromonas* spp.) enriched from an active MFC (Figure 2). Before comparing between processes governing microbial community assembly and succession on clean anodes to those pre-applied with EDC, three independent experiments (total N=16) were conducted to determine average startup rates of replicate MFC reactors inoculated using a fresh diluted wastewater (10% v/v) sample as the microbial source (wastewater was sampled the day prior to experiment initiation; therefore, each experiment (1-3) had a different starting microbial composition, Supplementary Figure S1). The development of MFCs’ current density was best described by four different phases (Supplementary Figure S1): **I.** Lag phase (designated as lag time, T_λ_): Base-line current production (0-9.2 mA/m^2^). **II**. Initial phase: initial current production (defined as > 9.2 mA/m^2^). **III**. Exponential phase: a rapid (close to linear) increase in current. **IV**. The mature phase: current production reaches an asymptotic value, representing the maximal current possible for this system when organic substrate is not limited. The lag phase is defined here as the time until the MFC reaches 9.2 mA/m^2^. This value was chosen arbitrarily as a value that is higher than base-line current, indicating that the biofilm is starting to acclimate to the anode, yet it is still in an initial stage of development. Startup time was defined as the time to reach steady-state current production, i.e. the time to reach the maximum current that a particular anodic biofilm is able to produce under specific given conditions.

### Applying the enriched Desulfuromonas Consortium on the anode surface

We specifically chose to sample an anode from an MFC that had been operating for 40-days (reached steady state voltage generation). Based on previous results (Yanuka-Golub et al., 2016), the anode was assumed to be colonized by members of the *Desulfuromonadaceae* family; and this was later verified by 16S rRNA gene amplicon sequencing (Supplementary Figure S2). Anode samples were plated on bicarbonate sodium fumarate plates and incubated anaerobically until light-orange-colored colonies appeared after three weeks (Badalamenti et al., 2016; Finster et al., 1997; Kuever et al., 2015). For taxonomic identification, the 16S rRNA gene of selected colonies was sequenced and compared to NCBI database. Notably, while the orange colonies were identified as belonging to the *Desulfuromonas* genus, we were not able to obtain them in an entirely pure culture, despite several rounds of isolation on solid media; we thus considered them as an enriched *Desulfuromonas* consortium, hereafter referred to as “**EDC**”. For each MFC, several EDC colonies collected from the same Petri dish were applied directly on the anode surface (i.e. bioaugmentation). To examine the microbial composition of the original consortium, EDC was sequenced via 16S rRNA amplicon sequencing and shotgun metagenomics. Both methods showed that *Desulfuromonas acetexigens* was the species most closely related to the *Desulfuromonas* enriched in EDC (this was verified by mapping shotgun metagenome reads to the *recA* gene that served as a taxonomic marker).

The consortium was directly applied onto a clean anode in MFCs inoculated either with a diluted wastewater sample (N = 5) or with a diluted filtered wastewater (N = 5), 10% v/v. Filtered wastewater was treated as following: raw wastewater was first filtered through Whatman® Grade 1 Qualitative Filtration Paper to remove bigger particles, and then through a 0.22 μm filter (Stericup-GP, 0.22 μm, polyethersulfone, Merck) for the removal of the majority of microorganisms present in the sample. Additionally, as a control group, MFCs with a clean anode (i.e. not applied with enriched electroactive-consortium) were inoculated with only a diluted wastewater sample as the inoculum source (N = 4). Additional anode samples were taken at different time points from bioreactors that initially contained only bicarbonate buffer and EDC augmented anodes (N=6). After approximately 42 hours, the voltage of all six reactors dropped (Supplementary Figure S3). At this time point (day ~1.75) the anode was sampled (referred to as EDC+B). Following, the media of one selected reactor was immediately switched to diluted wastewater, which resulted in an immediate voltage increase. After maximum current density was reached, the anode was sampled again (referred to as EDC+B--WW), Figure 2B. Essentially, EDC+B--WW resembled EDC+WW, experimentally, except EDC+WW was inoculated with diluted wastewater from day zero and EDC+B--WW after 42 hours.

Details of the how electroactive bacteria were enriched from a working MFC anode is described in Figure 2A and in Supplementary methods.

### DNA extraction and microbial community analysis

Anodic biofilms and planktonic cells from the MFC systems were sampled at different time points for community and phylogenetic analysis, as previously described (Yanuka-Golub et al., 2016). For treatment groups EDC+WW, EDC+FWW and WW, planktonic cells were sampled at three time points (day 1, day 3 and time at which steady-state current production was achieved – SS; N = 2). Day zero was considered as either filtered or unfiltered inoculum source. Briefly, total DNA was extracted from anodic biofilms or from the suspended cells using the PowerSoil DNA isolation kit (MO BIO Laboratories) according to the manufacturer’s instructions. For 16S rRNA amplicon, 72 samples were amplified using the 341F-806R primer set, sequenced and analyzed (Supplementary Methods). An OTU table was used as input for phyloseq (McMurdie and Holmes, 2013) and DESeq2 (Love et al., 2014) R packages. Bacterial diversity within samples (alpha diversity) was estimated using Richness and three diversity indices, Simpson, Shannon and Pielou’s evenness. In order to investigate the enrichment of specific populations in the planktonic or in the anode’s microbial community, we performed the DESeq function in the DESeq2 R package to contrast the anodic and planktonic microbial communities of WW and EDC+WW. DESeq2 uses negative binomial models and raw read counts to test if each OTU is differentially abundant across experimental factors. Using this function, we selected the OTUs that were enriched in at least one of the microbiomes. Metagenome assembly, gene prediction and functional profiles were completed for six DNA samples (6 out of the same 72 genomic DNA samples that were sequenced for 16S amplicon). Additionally, the putative genome of *D.acetexigens* was assembled from four samples and average nucleotide identity (ANI) was calculated. The methods used for DNA extraction, 16S rRNA gene and shotgun metagenome sequencing and library construction and metagenome assembly analysis are fully described in the Supplementary Methods.

### Electrochemical Measurements

Voltage was measured across a fixed external resistor (R ext = 1000 Ω). The current (I) was calculated according to Ohm’s law: I = E/R. For polarization curves, voltage was plotted as a function of current density (A m^−2^). A resistor box was used to set variable external loads while recording the voltage for each external resister applied, and the current was calculated using Ohm’s Law. The voltage value was recorded at fixed time gaps (10-15 minutes) after steady-state conditions have been established under the specific external resistance employed.

### Chemical Measurement

Water samples were filtered prior to the chemical analysis, unless stated otherwise (Amicon Ultra 0.5 ml 10K, Mercury LTD., Israel). The methods used for all chemical analyses of water samples can be found in the Supplementary Methods.

## Results

### Isolation of an enriched electroactive consortium and MFC bioaugmentation reduces time to stable current production

Previous results have shown that electroactive biofilm assembly process on clean anode surfaces, starting from the same inoculum can be highly stochastic, leading to different startup times and community composition (Yanuka-Golub et al., 2016). We repeated this on a larger scale, following sixteen reactors inoculated with diluted wastewater and clean anode surfaces for as long as it took them to achieve stable current production (defined as “start-up time”, Figure 1). Startup time again showed high variation between cells (18.83±7.11 days), while lag times (the time to achieve primary current production, see methods) tended to be more similar among the 16 reactors (4.92±1.04 days, Figure 1).

**Figure 1:**
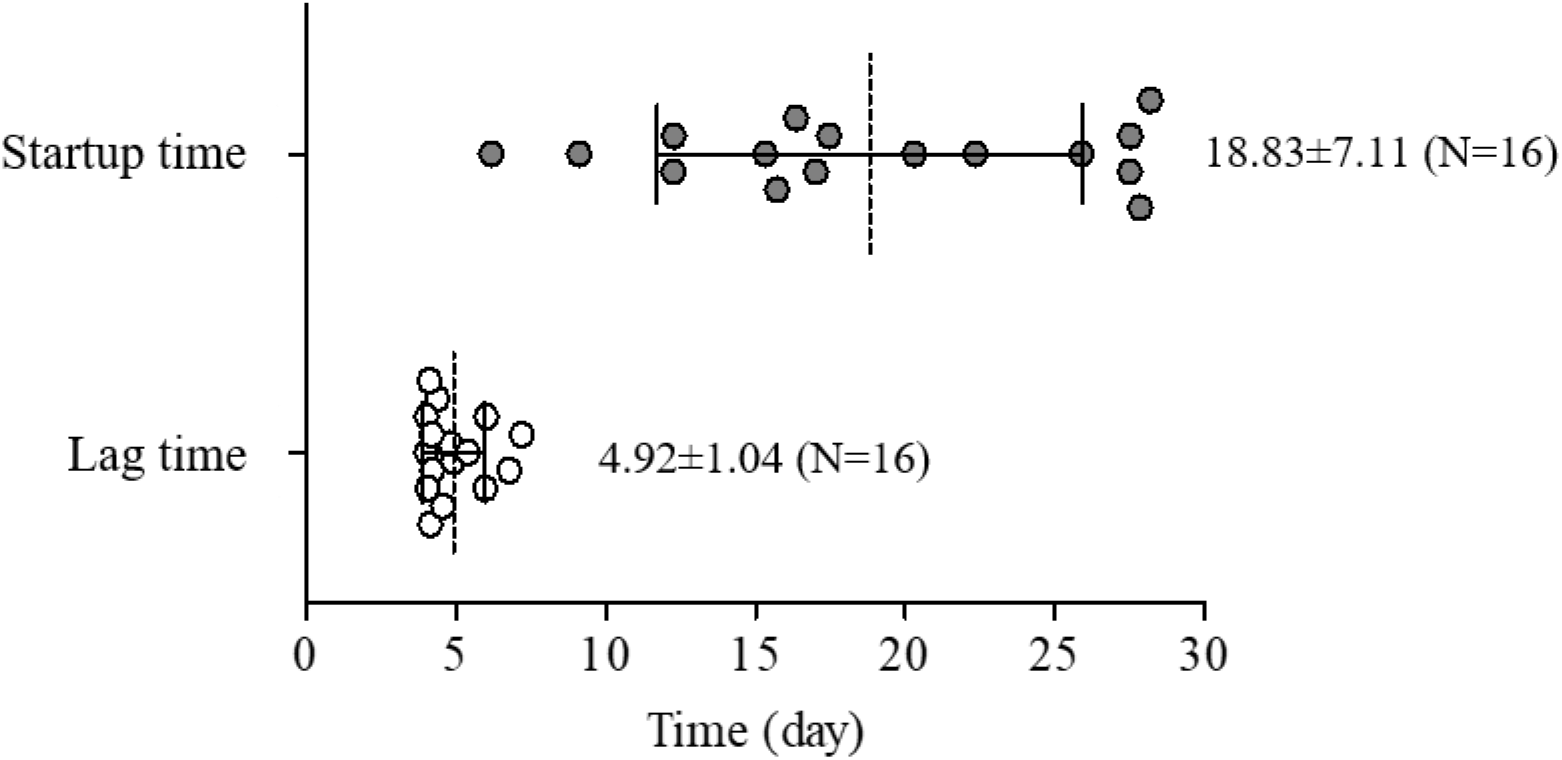
Scatter dot plot comparing lag time and startup time for replicate MFC reactors (N =16) inoculated with diluted wastewater as the microbial source from three independent experiments. The horizontal line indicates the mean with error bars indicating standard deviation.

We hypothesized that in order to reduce the high stochasticity characterizing early biofilm assembly, deterministic factors should be enhanced by pre-colonizing the anode surface with a defined microbial community composed primarily of electroactive species. We thus augmented MFC anodes by directly applying a microbial electroactive consortium isolated from a working anode (Figure 2A). This consortium (EDC) was dominated by two or more *Desulfuromonas* strains that were distantly related to *D. acetexigens* (ANI of 91.5-92.1%, Figure 3), but relatively similar to one another (ANI of 99-99.4%) in all EDC batches, indicating that similar strains of *Desulfuromonas* colonized all replicate anode surfaces (Figure 3).

**Figure 2:**
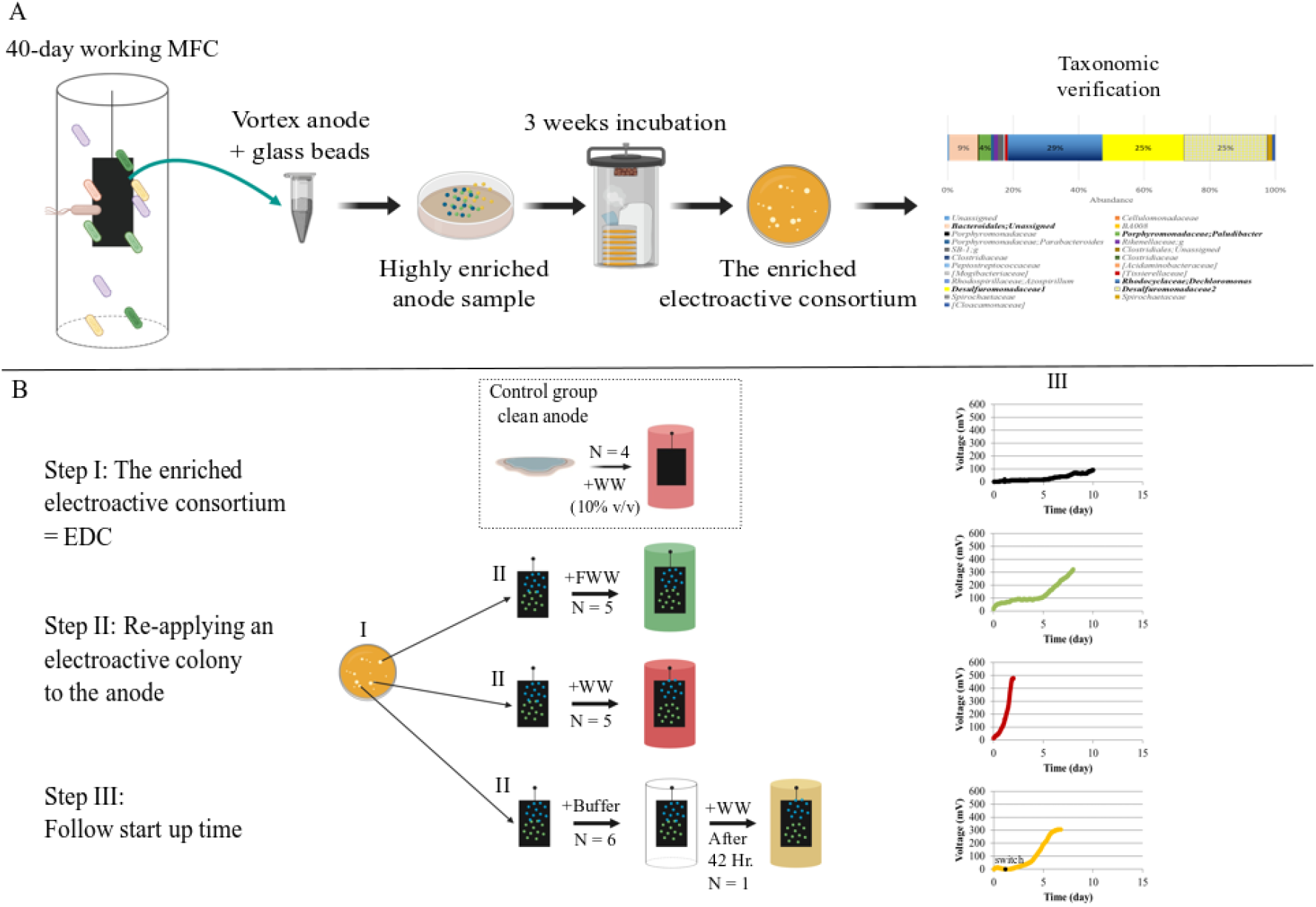
Schematic representation of experimental design and treatment groups. A) Working scheme showing the way the electroactive enrichment (E**DC**) was obtained from an anode sample taken from a 40-day working MFC, incubated for several weeks, resulting in growth of colonies that were taxonomically verified by 16S**-**amplicon sequencing to contain electroactive organisms. B) Schematic representation of the experimental design.

**Figure 3:**
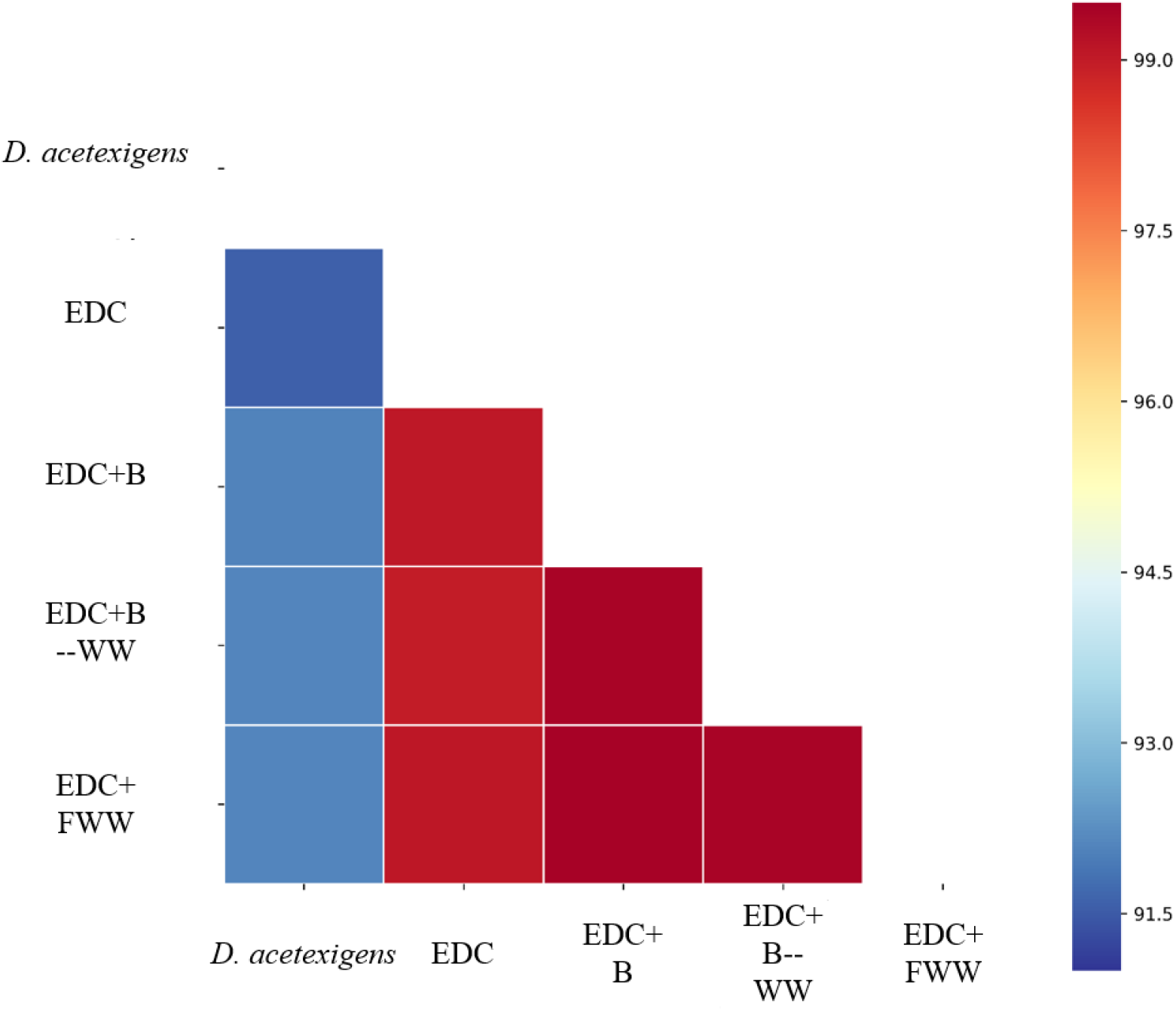
Heatmap of pairwise average nucleotide identity (ANI) values for the genome of *D.acetexigens* that was separately assembled from the four samples presented. *D.acetexigens* (Katuri et al., 2017) was used as a reference genome.

When anodes to which EDC was applied were placed in an MFC containing bicarbonate buffer and acetate as a carbon source (N = 6), voltage increased rapidly, but quickly dropped to zero shortly after (Supplementary Figure S3A-F). Recovery of current was observed when the MFC’s bulk liquid was switched to either dilute wastewater or dilute filtered wastewater (10% v/v) immediately after that drop (Supplementary Figure S3B-F). To investigate this phenomenon, three different treatments were compared, as illustrated in Figure 2B. The first group of MFCs (N = 5) was inoculated with diluted filtered wastewater (EDC+FWW), and a second group (N = 5) was inoculated similarly with diluted wastewater including their entire microbial diversity (EDC+WW). The anode surfaces of both these groups were pre-treated with EDC, as described above. A control group of MFCs (N = 4) was inoculated with wastewater without applying EDC to the anode (i.e. a clean anode surface). All solutions were supplemented with acetate. This experiment demonstrated that both lag time and startup time were significantly lower in bioreactors that had the EDC-augmented anodes with diluted wastewater, compared to control MFCs (Figure 4A and 4B; Kruskal-Wallis test; p<0.05). Notably, mean lag time and startup times of EDC-augmented anodes with filtered wastewater showed a moderate improvement. The filtration treatment did not affect the chemical composition of essential components in wastewater, with the exception of organic nitrogen, which was two-fold reduced in filtered wastewater (Supplementary Figure S4). 16S rRNA gene amplicon sequencing showed that, as expected, filtered wastewater used for inoculation (i.e. day 0) lacked many of the bacteria present in unfiltered wastewater, and consequently has lower diversity (Shannon of 6.2±0.2 vs. 9.7±0.03 and Faith’s phylogenetic diversity (PD) of 10.1±2.02 vs. 59.5±1.6, respectively). The filtered wastewater community comprised of several small cell taxa that could potentially pass through the 0.22 µm filter (Supplementary Figure S5 & S6). To examine the contribution of these species to the electroactive biofilm formation, a different control MFC group, inoculated with filtered wastewater with a clean anode (henceforth blank control) was tested, and showed a slight increase in voltage over time (Supplementary Figure S7), indicating that this inoculum has some limited electrogenic biofilm forming potential. In contrast, no difference was observed among the three treatments for maximal current density obtained (Figure 4C). Therefore, we next asked which factor(s) led to such a strong synergy between the wastewater "source community" composition and the pre-colonized anode community, which together achieved a dramatically higher functional anode biofilm.

**Figure 4:**
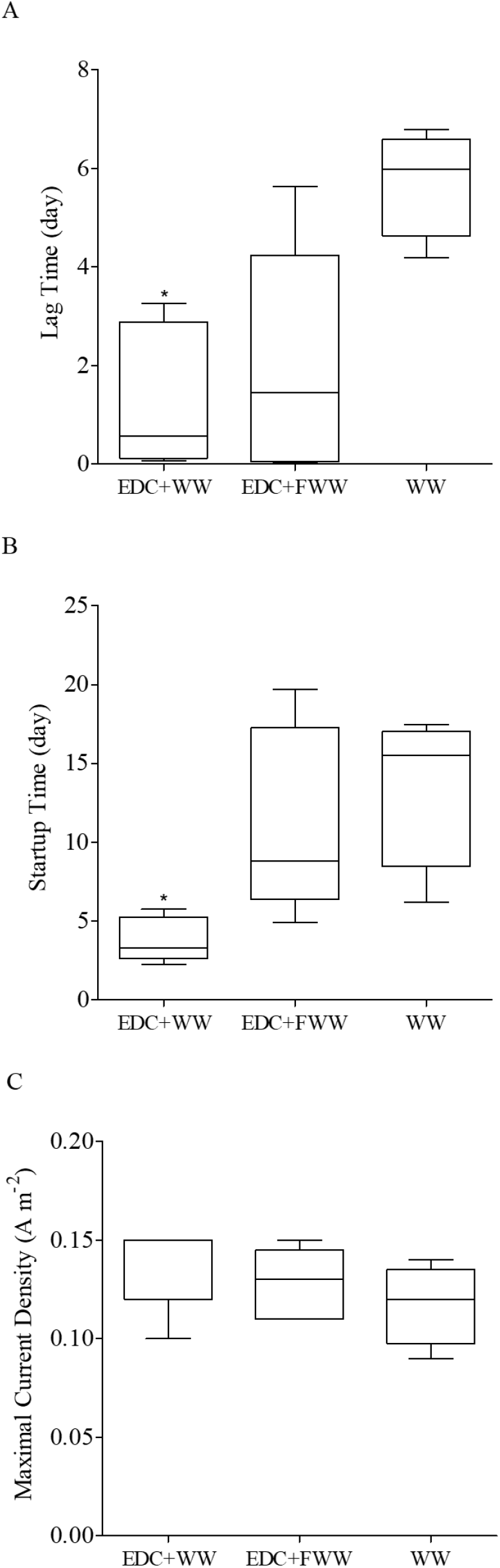
A-C) Startup time, lag time and maximal current density of the MFC reactors of three treatment groups: EDC+WW, EDC+FWW and WW, respectively.

### Predictable development of a highly-functional anode-colonizing biofilm requires specific water-borne bacteria

Potentially, augmentation of the anode with an electroactive consortium should not only shorten startup time but also result in more predictable MFC communities, similar to one another. To compare the microbial community composition of each of the pre-colonized anode surface (EDC+FWW and EDC+WW treatment groups) to those of non-pretreated anodes (WW), the anodic microbial communities of all reactors were collected and compared once stable current density was obtained (defined as steady-state). Additional anode samples were taken at different time points from bioreactors that initially contained only bicarbonate buffer with acetate and EDC augmented anodes (N=6), after the voltage drop (see above) at 42 h., some of which were supplemented with wastewater (see methods, and figure 2B). The planktonic community for each of the treatment groups was also sampled at different time points (total 48 samples, see methods). The importance of analyzing the planktonic community dynamics lies in the two potential sources of the planktonic bacteria. The first is the wastewater in the MFC that provide a microbial source (i.e. a pool of newly-arriving species), and the second is cells that are released from the anodic biofilm into the bulk liquid. The latter implies that the planktonic community can potentially reflect the anodic biofilm assembly (Yanuka-Golub et al., 2016), without harming the anode. Following planktonic community dynamics is especially important during the very early stages when the interaction between deterministic and stochastic processes can substantially affect the interaction between the new-arriving species with the founders (Tilman, 2004). All microbial communities were analyzed by 16S rRNA gene amplicon sequencing (see Methods).

Overall the communities could be assigned to four clusters according to their composition (PcoA analysis, Figure 5A-B) regardless of the distance measure used (Bray-Curtis, weighted or unweighted UniFrac). The first cluster contained samples that were similar in taxonomic composition to the EDC - EDC+B, EDC+FWW and EDC+B—WW (henceforth ***cluster I***). The second cluster comprised of samples that had a statistically significant positive effect on both lag time and startup time (Figure 4A-B) relative to non-bioaugmented MFCs (EDC+WW, henceforth ***cluster II***). The third group included clean anodes that were not pretreated with EDC (i.e. WW; ***cluster III***), and the fourth cluster comprised of blank controls (MFCs inoculated with filtered wastewater and a clean anode). Analysis of similarity (ANOSIM) of anode communities indicated that cluster I was significantly different from clusters II and III, but the clusters differed to a larger degree in the planktonic compared to the anodic communities (Supplementary Table 4). On the other hand, while anode samples of cluster II and III were somewhat similar according to the weighted distances, the planktonic samples of these two groups clustered exclusively together according to all distance measures (Figure 5B, Supplementary Table 4). This may suggest that the difference between cluster I and clusters II/III was caused by the emergence of specific taxonomic groups originating from the wastewater “source community”, as well as specific interspecies interactions that may have occurred between those “source communities” and the communities that already pre-colonized the anode by applying EDC on its surface.

**Figure 5:**
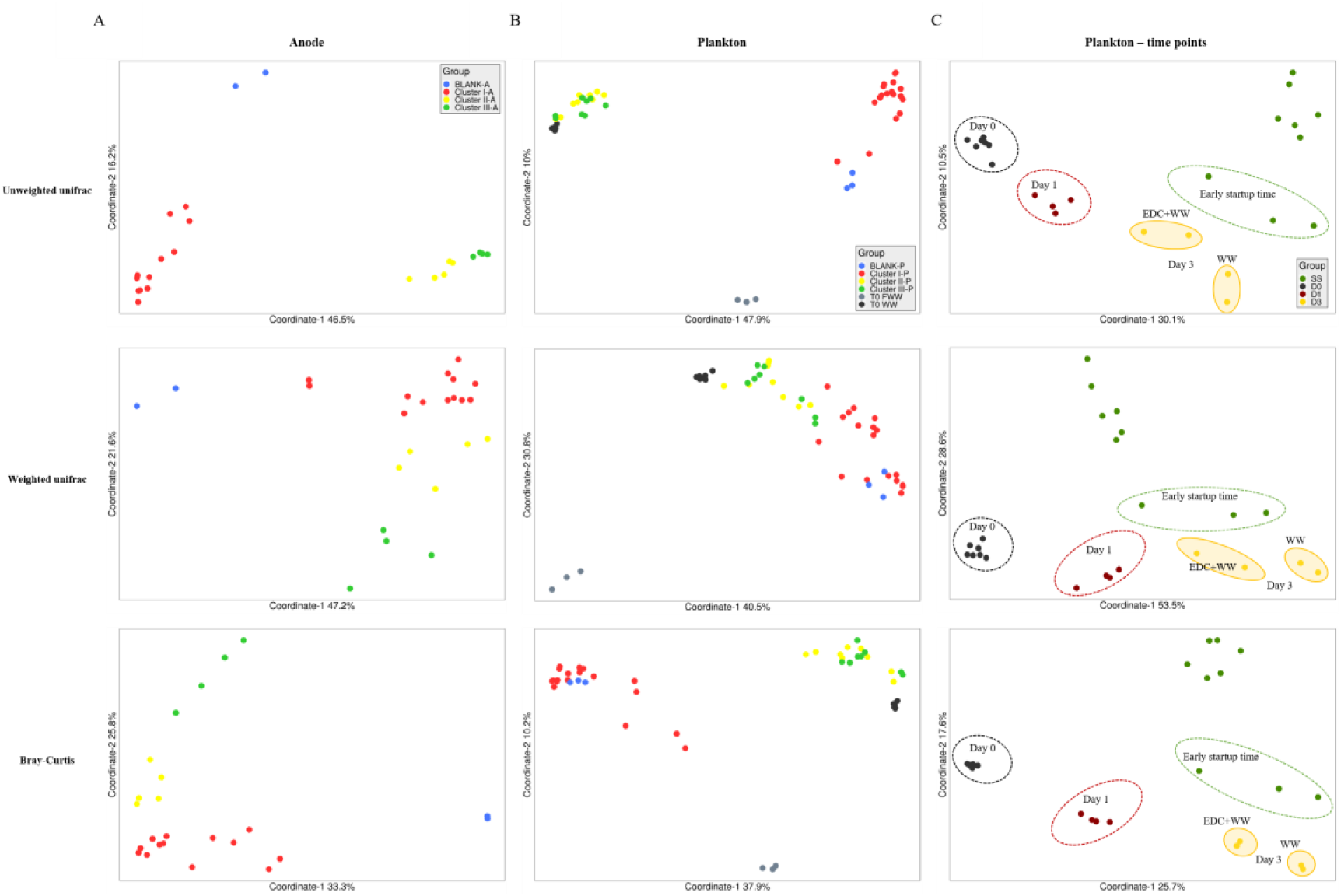
Principal Coordinate Analysis (OTU level) based on weighted and unweighted Unifrac, and Bray-Curis distances of the A) anode-biofilm communities: BLANK: control of filtered wastewater with a clean anode, Cluster I: EDC, EDC+B, EDC+FWW, EDC+B—WW, Cluster II: EDC+WW, Cluster III: control – MFC supplemented with wastewater (WW) and clean anode B) plankton communities and C) plankton communities of only clusters II and III at specific time points: day 0 at which wastewater was supplemented with the bioaugmented anode, D1: day 1 after inoculation, D3: day 3, SS: Steady-State – the time at which stable maximal current density was achieved. This day varied between each MFC.

It was intriguing that although both anodic and planktonic communities of cluster II and cluster III (EDC+WW and WW, respectively) were taxonomically similar, each system exhibited a substantial different startup time (Figure 4A-B). While alpha diversity indices, except PD (Shannon and Simpson) of the clean anode samples were significantly higher than those of the EDC pre-colonized anode surface, all three alpha diversity indices’ values of the total planktonic community did not differ between EDC+WW and WW groups (Figure 6A). Nevertheless, already on the first day (day 1), PD differed between the two groups, while no differences were observed for other alpha (Figure 6B) and beta diversity distances (Figure 5C) at this time point. Later, on day 3, the samples of both treatment groups clustered separately (Figure 5C), in agreement with the three alpha diversity indices at this time point (Figure 6B). Remarkably, the mean lag time of EDC+WW was 1.3 (±1.5) days and the time it took the bioreactors to reach maximal current density was between 2.2 and 5.7 days (3.8±1.4 days, Figure 4B), coinciding with the observed community divergence on day 1 and 3, respectively. Overall, these results indicate that specific planktonic community assembly processes occurred at very early time points in MFCs pre-treated with EDC, which subsequently affected their corresponding anodic community structure and startup time. Finally, once steady state conditions were achieved, the planktonic communities of both treatments converged (Figure 6B). To explain the observed differences between EDC+WW and WW samples, DESeq2 was used to identify differentially abundant amplicon sequences in EDC+WW group (positive fold change) compared to WW group (negative fold change indicate OTUs enriched in WW). Anode and planktonic samples were analyzed separately. When we compared anode samples of EDC+WW to WW samples, DESeq2 identified 49 sequences that were significantly differentially abundant in EDC+WW (Figure 7A). Comparing the planktonic sequences of day 1 showed that only one OTU was significantly differentially abundant in EDC+WW (from *Flavobacteriia*), subsequently, this OTU on day 3 became differentially abundant in the WW planktonic community, and was replaced by a different OTU (from *Sphingobacteriia*) in the EDC+WW planktonic community (Figure 7B). In steady-state, 63 sequences were identified to be significantly differentially abundant in the WW planktonic community (Figure 7B), in which Alphaproteobacteria, Clostridia and Flavobacteriia were the most abundant groups.

**Figure 6:**
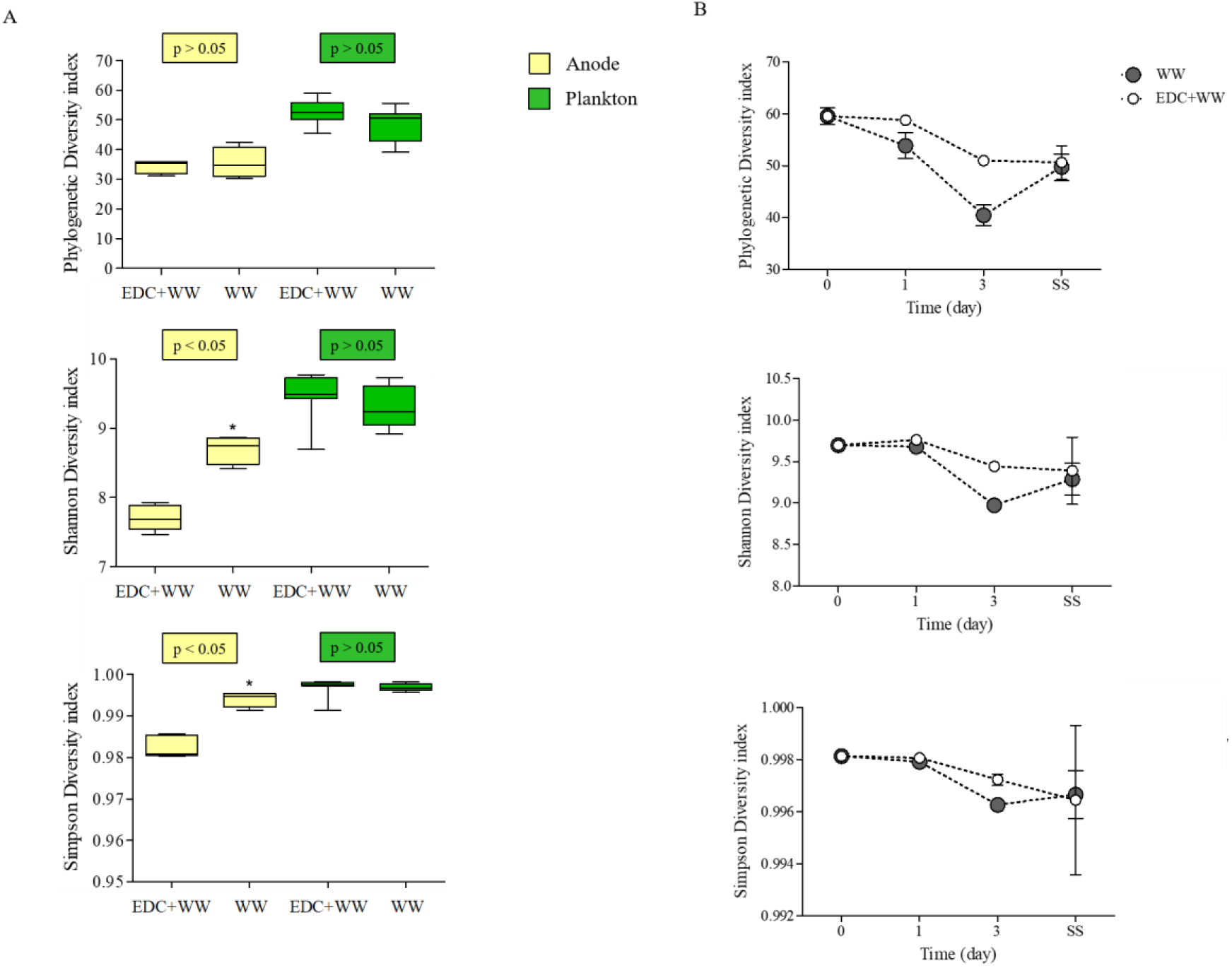
A) Alpha diversity for anode-attached communities and planktonic communities in the MFCs of two groups: augmented anodes (EDC+WW) and control non-augmented anodes (WW). B) Temporal dynamics of the planktonic communities of the same two groups. The last time point is SS, indicating the time at which steady-state current production was achieved. For the WW group this time was significantly longer than EDC+WW as explained in text.

**Figure 7:**
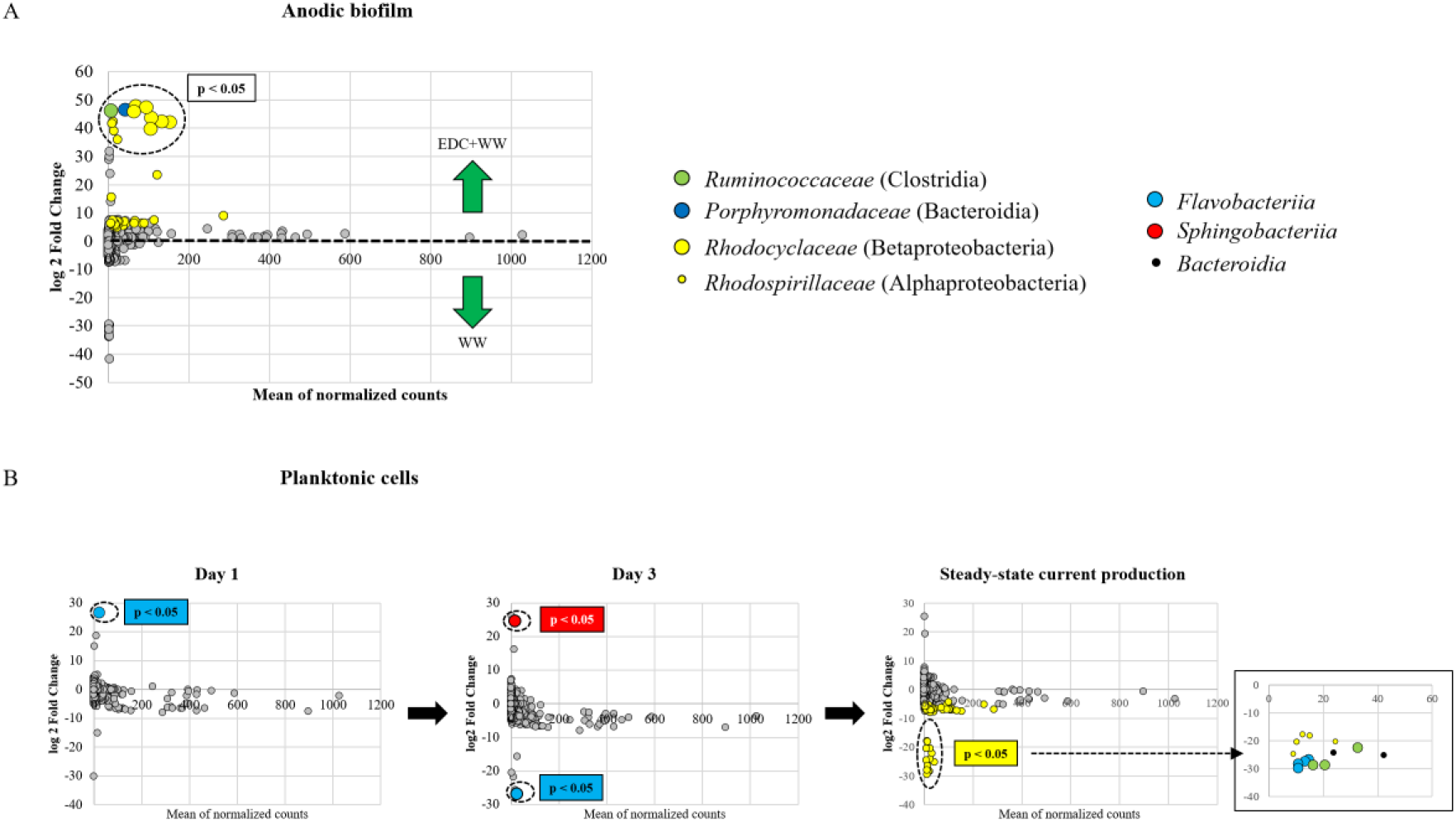
DESeq2 analysis results indicating the fold-change of OTUs of MFCs that their anode-surface was bioaugmented relative to control non-bioaugmened anode surfaces (p < 0.05) in A) anode-attached communities and B) Planktonic communities.

To obtain a higher taxonomic resolution of the established anodic biofilm and the differences between clusters I-III, as well as functional potential of the genes that they encode, six representative anode samples were sequenced via shotgun metagenomics and the obtained reads were assigned taxonomic and functional classifications. Indeed, metagenomics data of the six representative anode samples was in agreement with the 16S amplicon sequencing with respect to the most differentially abundant taxonomic groups identified according to DESeq analysis (Figure 7A), as well as, the strong taxonomic association between the EDC+WW and WW anode samples (Supplementary Figure S8). Taxonomic observations of shotgun metagenomics data revealed important differences between cluster I and II, Figure 8A. First, the most abundant family belonging to Deltaproteobacteria in cluster I was *Desulfuromonadaceae*, originating from the bioaugmenting consortium, as described above. However, the most abundant family in cluster II was *Geobacteraceae*, which reached 39% of total community. While in sample EDC+WW *Geobacter lovleyi* was the single species that dominated (31%), in WW without an EDC pretreatment, other *Geobacteraceae* species were more abundant relative to other samples. *Azonexus hydrophilus* was the most dominant species in cluster I aside from members of the *Desulfuromonadaceae*. In samples WW, EDC+WW, and EDC+B--WW, all of which had wastewater supplementation, relative abundance of *A. hydrophilus* was <1%. The relative abundance of Gammaproteobacteria was the highest in sample EDC+B--WW, in which *Pseudomonas stutzeri* was dominant. On the other hand, *Shewanella* spp., a well-known electroactive organism belonging to Gammaproteobacteria was mainly enriched in sample EDC+WW.

**Figure 8:**
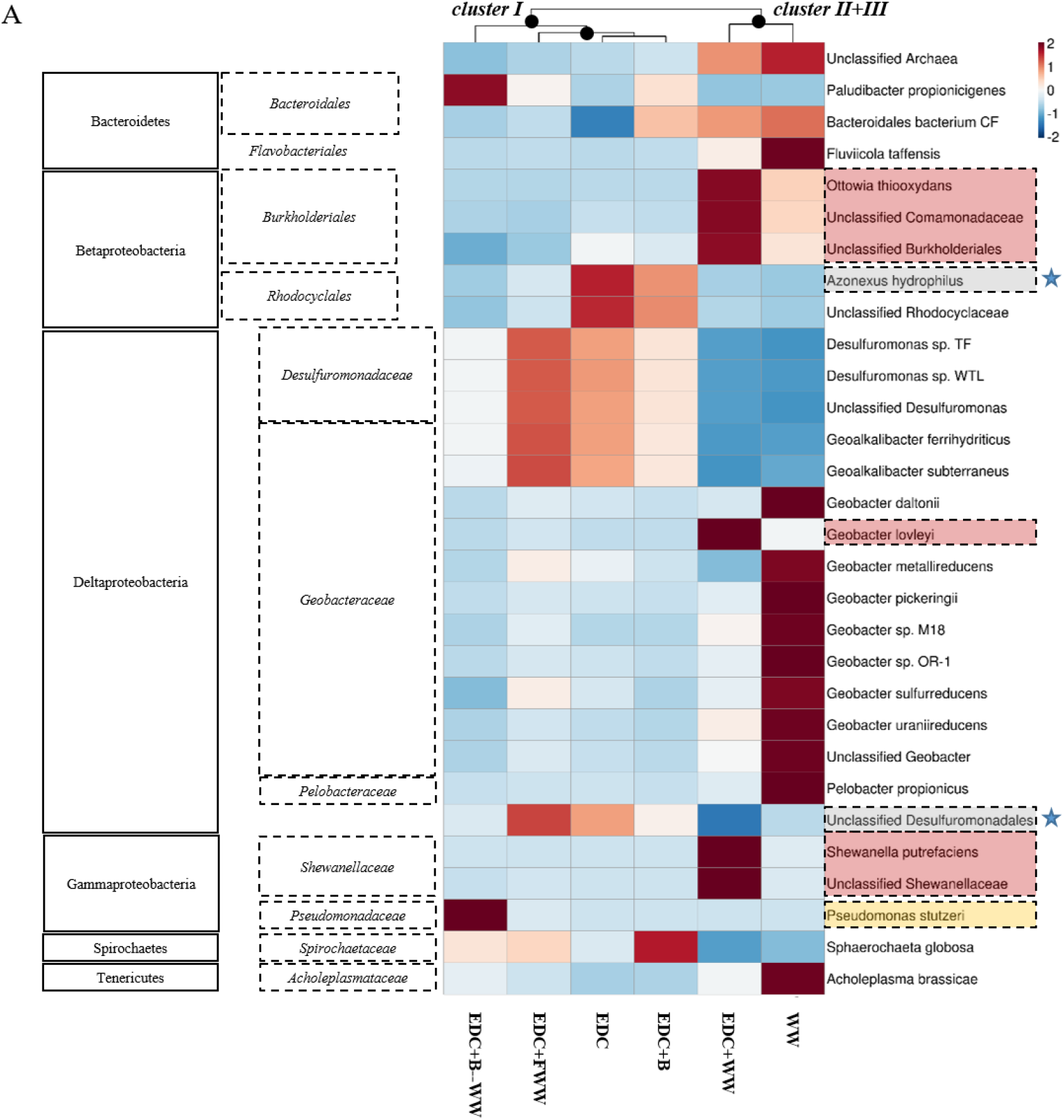
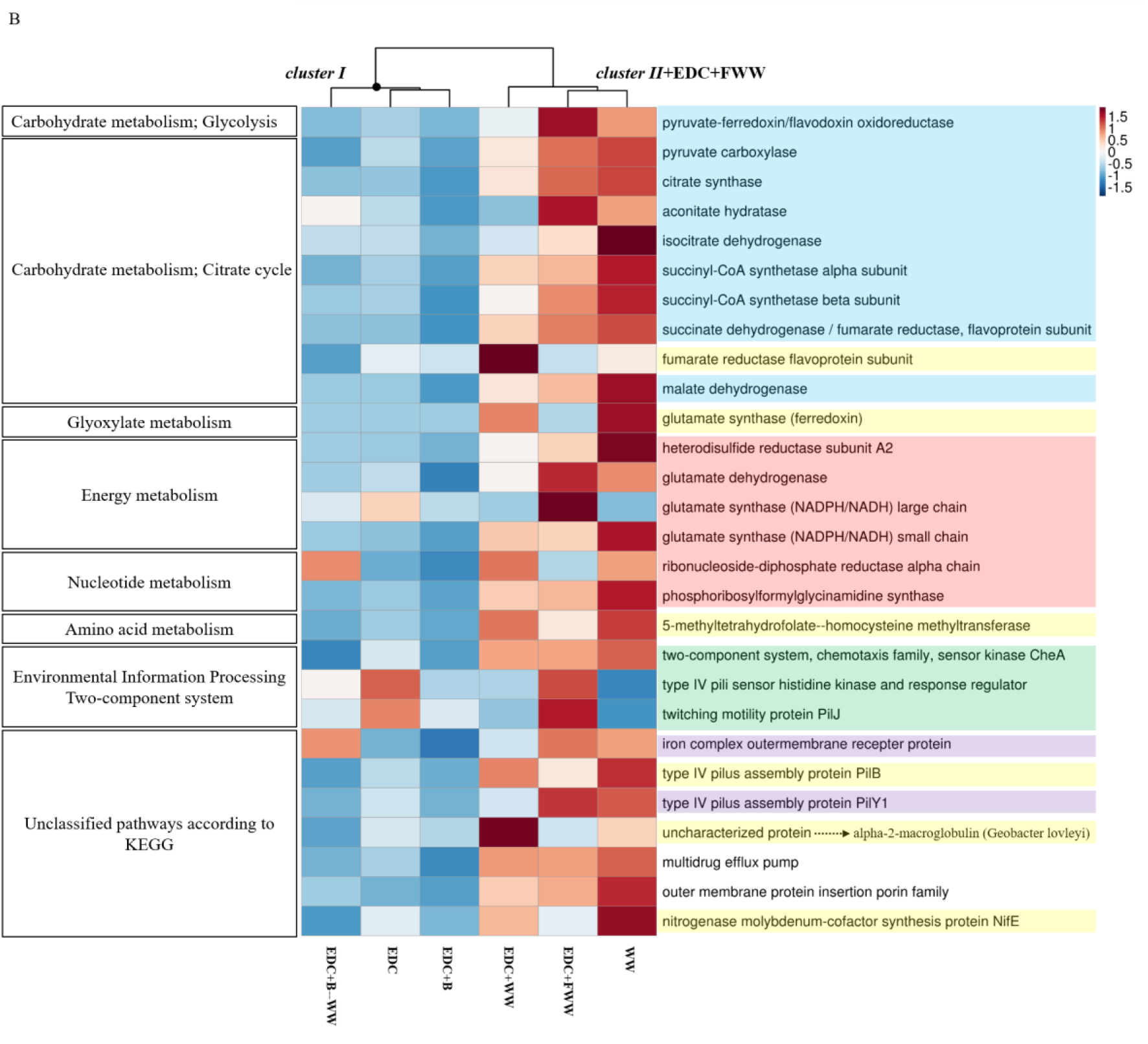
Heatmaps of A) bacterial taxonomic composition. B) KEGG ortholog functions. Both heatmaps are based on shotgun metagenomics and clustering of the samples was computed using Bray-Curtis dissimilarity index.

Comparing anode samples applied with EDC inoculated with clean buffer to those inoculated with filtered wastewater revealed potential essential bacteria that were lacking in EDC, including members of *Flavobacteriales*, *Sphingobacteriales*, *Rhizobiales* and *Rhodospirillales*. Interestingly, amplicon sequences belonging to these classes were detected by DESeq2 as differential when compared between groups WW and EDC+WW (Figure 7A), further supporting their importance as key members during biofilm formation.

### The highly-functional anode-colonizing community was linked to chemotaxis regulation, type IV pilus assembly and colonization proteins

To better understand how the taxonomic differences correspond to functional differences, we compared the abundances of KEGG ortholog functions (KOs) in the six anode biofilm communities. Overall, the six samples had a relatively similar functional profile with two clusters splitting at ~75% similarity (based on Bray-Curtis dissimilarity index, Supplementary Figure S8). Notably, while the anode sample EDC+FWW was related in terms of predicted functions to samples WW and EDC+WW, taxonomically it clustered with EDC and EDC+B (Figure 8A and 8B). A higher number of proteins associated with motility, chemotaxis, signal transduction, membrane transport and carbohydrate metabolism was observed in the anodic biofilms belonging to cluster II and sample EDC+FWW. Sample EDC+WW had a unique functional profile, explicitly resulting from the coupling of wastewater and the pre-colonized anode (Figure 8B). Six protein coding genes were found to be specifically associated with samples EDC+WW and WW and not in EDC+FWW (marked yellow, Figure 8B). Four were associated with KEGG metabolic pathways (fumarate reductase, glutamate synthase, nifE and homocysteine methyltransferase). Fumarate reductase, an important enzyme for anaerobic respiration (Galushko and Schink, 2000), was primarily encoded by *A. hydrophilus* in EDC and *G. lovleyi* in EDC+WW, and had the highest abundance in EDC+WW. This probably indicates the specific contribution of *G. lovleyi* to acetate oxidation through central metabolic pathways, leading to an efficient EET that eventually benefits the entire anodic biofilm. Other metabolic genes that were *Geobacter*-associated were glutamate synthase, an iron–sulfur flavoprotein that plays a key role in ammonia assimilation (Vanoni and Curti, 1999) and 5-methyltetrahydrofolate--homocysteine methyltransferase, previously detected as one of the most highly expressed genes on closed circuit electrode surfaces (Marshall et al., 2017). The nitrogenase molybdenum-cofactor biosynthesis protein NifE has a critical role in nitrogen fixation (Brigle et al., 1987). NifE was primarily encoded by *Geobacter* in EDC+WW and WW anode samples and by *Desulfuromonas* and *A. hydrophilus* in EDC, EDC+B and EDC+FWW. Additional differential functions were type IV pilus assembly protein PilB and an uncharacterized protein homologous to alpha-2-macroglobulin of *G. lovleyi*, which are likely to be important in surface colonization (Budd et al., 2004; Steidl et al., 2016).

## Discussion

Random and non-random processes that govern microbial community assembly and succession on surfaces have long been of great interest for a wide range of community ecology disciplines, including plants, algae and microorganisms (Jackson, 2003). In microbial systems, most studies are associated particularly with the colonization of new substrates, following the model suggested by Jackson (2003), which describes the changes in community properties that occur during bacterial biofilm succession starting from the very early stages until the biofilm is well established. Conversely, in this study, we explored which factors drive biofilm assembly on a pre-colonized surface, which is especially relevant when applying bioaugmentation for *in-situ* bioremediation of natural environments (Tyagi et al., 2011) or wastewater treatment (Herrero and Stuckey, 2015). In groundwater, it has been shown that the composition of sediment-attached communities can differ substantially compared to suspended planktonic communities (Fillinger et al., 2019; Lehman et al., 2001), therefore, using MFC reactors as a model system allows full control of the environmental conditions and easily differentiating between surface-attached (i.e. anodic biofilm) and suspended communities (Zhang et al., 2019; Zhou et al., 2013). Furthermore, MFCs’ current density serves as a bio-signal directly associated with the anodic microbial community performance and biofilm formation stages with time, as illustrated in Figure 1 and 3.

Direct application of an enriched consortium combined with wastewater substantially reduced startup times to 3.8 days from an average of 18 days observed for sixteen empty anode surface. Interestingly, the immediate voltage produced was not stable unless the anode surface was bioaugmented and also co-inoculated with either filtered wastewater or wastewater. Filtered wastewater had a reduced effect on startup time relative to non-filtered wastewater (Figure 4), in agreement with a recent study that found that the performance of bioelectrochemical reactors declined as biofilm diversity decreased (Zhang et al., 2019). The taxonomic composition of filtered wastewater anodes resembled EDC, and was substantially different from non-filtered wastewater anodes (Figures 5 and 8A). This implies that a successful anode-colonization process (of sample EDC+WW), which led to an extremely fast and robust startup time, involved a large shift in the anode-associated founder community. A particularly interesting switch was observed between the two main exo-electrogenic species, *Desulfuromonas* spp. and *Geobacter lovleyi*, which suggests a specific interaction that could exist between these closely related species in other systems, for example, sediment-attached communities in subsurface environments. Furthermore, the fact that this taxonomic shift did not occur in samples EDC+B, EDC+FWW and EDC+B--WW indicates that these samples had a relatively similar anode biofilm composition, further supporting the importance of the addition of the complete microbial source (wastewater instead of filtered wastewater) to rapidly achieve a high-functioning electroactive community on the anode surface. Since *Geobacteraceae* and *Desulfuromonadaceae* are phylogenetically related (Lonergan et al., 1996), occupy similar niches and use the same carbon sources and electron acceptors (both in engineered and natural systems), it is reasonable to hypothesize that when seeded at time zero, these groups (as planktonic, free-living cells in the bulk liquid) compete for the same resources (electron donors and available space to attach to the anode, colonize and transfer electrons to it).

Anode biofilm composition of the two groups EDC+WW and WW at steady-state conditions was highly similar, yet, MFCs exhibited substantial different startup times. This fundamental difference was hypothesized to occur already at early time points due to specific interactions between the planktonic community and EDC-pre-colonizers on the anode surface of EDC+WW. Further analysis revealed differentially abundant OTUs belonging to *Flavobacteriia* and *Sphingobacteriia* on day 1 and 3, respectively. On day 3, corresponding with the average startup time of EDC+WW reactors, the only OTU that became differentially abundant in EDC+WW relative to the control planktonic communities belonged to *Sphingobacteriia*. This finding is consistent with the functional role attributed to *Flavobacteriia* and *Sphingobacteriia*, both of which were found to represent key members in the formation and functioning of biofilms in diverse environments (Battin et al., 2016; Nagaraj et al., 2017; Pollet et al., 2018), probably due to their ability to degrade diverse complex organic material and their superior attachment ability. Furthermore, *Sphingobacteriia* were previously reported in bioelectrochemical devices both at the plankton fraction and anodic biofilm as homoacetogens or fermenters of complex organic substances that support electrogenic species within the biofilm through syntrophic interactions (Gao et al., 2014). Additionally, nitrogen fixation-related genes (*nifE* and *nifH*) were mostly abundant with EDC, EDC+WW and WW anode biofilms. While *nifE* in the latter two groups was affiliated with *Geobacteraceae* family members, in the former ones it was associated with *A, hydrophilus* and *D. acetexigens* (Supplementary Figure S9). Overall, the combined results from amplicon sequencing with the taxonomic and functional classifications obtained by shotgun metagenomics illustrate that only when supplemented with a full diversity inoculum of wastewater, a replacement of the founder species, EDC, (as well as key functional traits) on the anode biofilm occurred (i.e. a nearly complete turnover). Interestingly, even though the filtered wastewater did contain specific species that allowed an improved startup time relative to control MFCs, they did not eventually replace any of the founder species of EDC at steady state.

The question which ecological processes underlie the fate of founder species and to which degree their early abundance relate to their persistence as the community matures is still open and unexplored (Brislawn et al., 2019). Turnover is defined as an ecological process in which new taxa from a regional species pool are introduced into a given spatial domain; and successfully colonize it resulting in replacement of the existing taxa (Baselga, 2012). Alternatively, nestedness occurs when migration from the regional species pool is unsuccessful, leading to species loss without replacement of the invaded community by new arriving species (Baselga, 2010). Turnover and nestedness in turn were shown to influence microbial community dynamics and function in complex ecosystems (Graham et al., 2017; Stegen et al., 2018). However, we are still far from understanding comprehensively the feedback mechanisms that control these succession patterns, consequently limiting our ability to predict them. In this study, two different mature communities had very similar taxonomic and functional configurations; yet, the early-stage assembly processes substantially differed between them, resulting in different start up times. Thus, we show for the first time that pre-colonizing the anode surface with a defined electroactive consortium led to a strong synergy between the planktonic "source community" and the pre-colonized anode community. Essentially, we suggest that the pre-colonized anode created specific niches that became available and were quickly occupied by the newly-arriving species, especially key members that are biofilm formers (i.e. *Flavobacteriia* and *Sphingobacteriia*) that aided the electrogenic bacteria in establishing a highly-functional anode biofilm via syntrophic interactions. This enhanced niche-based processes dramatically stimulated the MFC performance, and thus offer a new approach for minimizing the system’s stochasticity and a better way to select for the most efficient microorganisms for environmental biotechnologies.

## Acknowledgements

The authors are grateful for the financial supports provided by the Israel Ministry of Environmental Protection (MOEP) under project agreement number 132-4-2 and the Tel-Aviv University Smaller-Winnikow Fellowship. In addition, we would like to personally thank Klimentiy Levkov for helping with the MFCs setup, and Stefan J. Green from the University of Illinois at Chicago DNA sequencing center for 16S amplicon sequencing.

