## Supplementary Figure S1 for "Anode surface bioaugmentation enhances deterministic biofilm assembly in microbial fuel cells"

### Slide 1
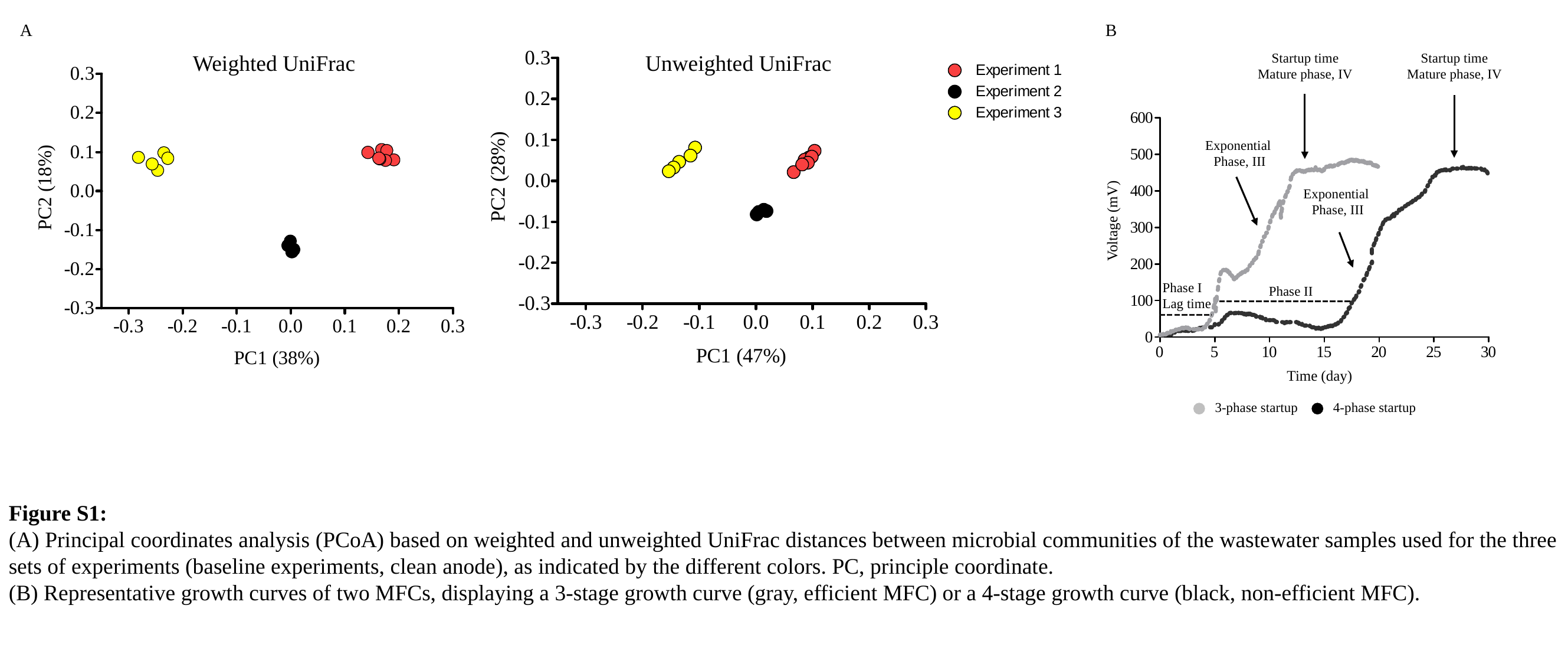

A
B
Weighted UniFrac
Unweighted UniFrac
Startup time
Mature phase, IV
Startup time
Mature phase, IV
Exponential
Phase, III
Exponential
Phase, III
Phase I
Lag time
Phase II
3-phase startup
4-phase startup
Voltage (mV)
Time (day)
Figure S1:
(A) Principal coordinates analysis (PCoA) based on weighted and unweighted UniFrac distances between microbial communities of the wastewater samples used for the three sets of experiments (baseline experiments, clean anode), as indicated by the different colors. PC, principle coordinate.
(B) Representative growth curves of two MFCs, displaying a 3-stage growth curve (gray, efficient MFC) or a 4-stage growth curve (black, non-efficient MFC).
