## Supplementary Figure S2 for "Anode surface bioaugmentation enhances deterministic biofilm assembly in microbial fuel cells"

### Slide 1
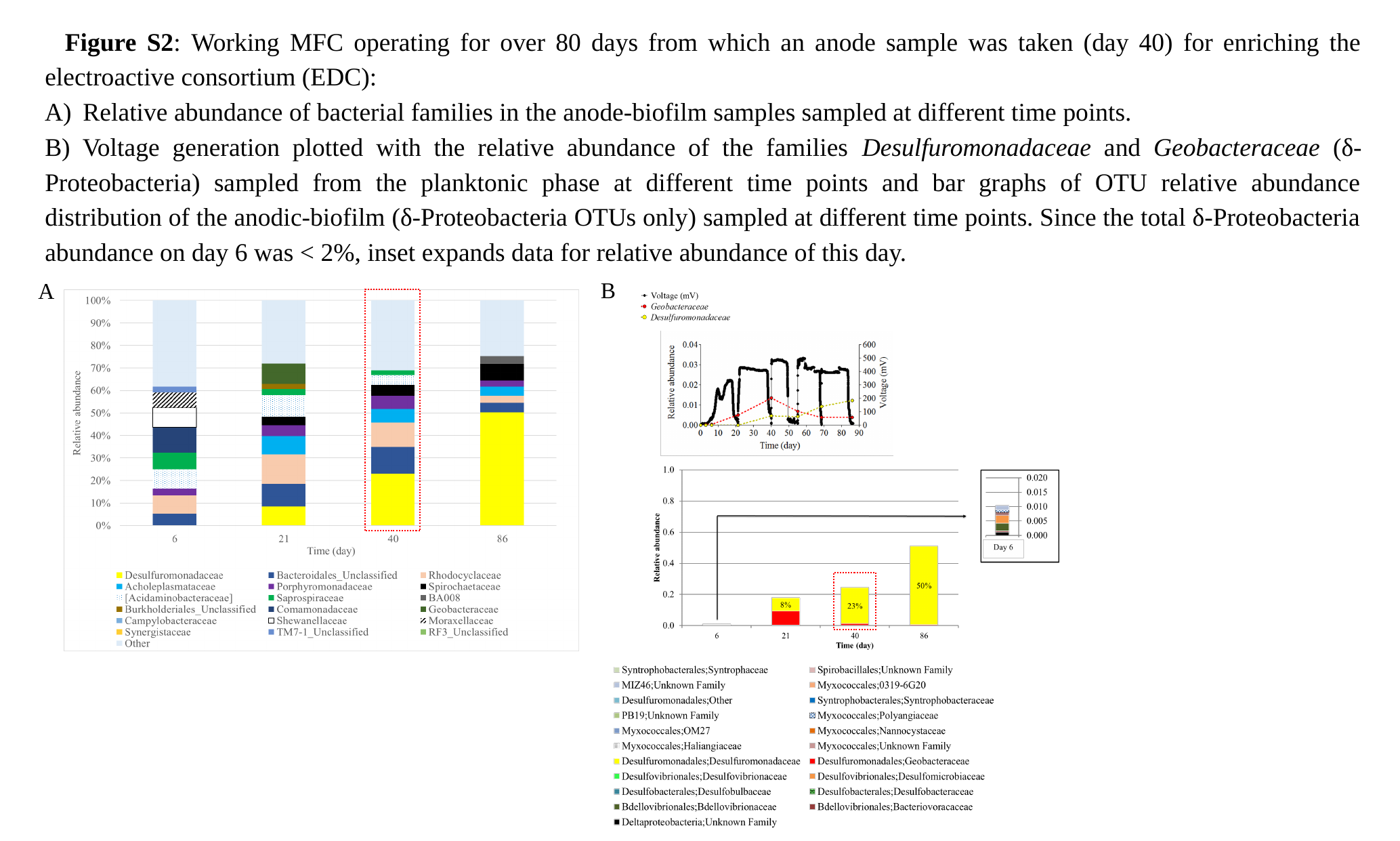

Figure S2: Working MFC operating for over 80 days from which an anode sample was taken (day 40) for enriching the electroactive consortium (EDC):
Relative abundance of bacterial families in the anode-biofilm samples sampled at different time points.
B) Voltage generation plotted with the relative abundance of the families Desulfuromonadaceae and Geobacteraceae (δ-Proteobacteria) sampled from the planktonic phase at different time points and bar graphs of OTU relative abundance distribution of the anodic-biofilm (δ-Proteobacteria OTUs only) sampled at different time points. Since the total δ-Proteobacteria
abundance on day 6 was < 2%, inset expands data for relative abundance of this day.
B
A
