## Supplementary Figure S3 for "Anode surface bioaugmentation enhances deterministic biofilm assembly in microbial fuel cells"

### Slide 1
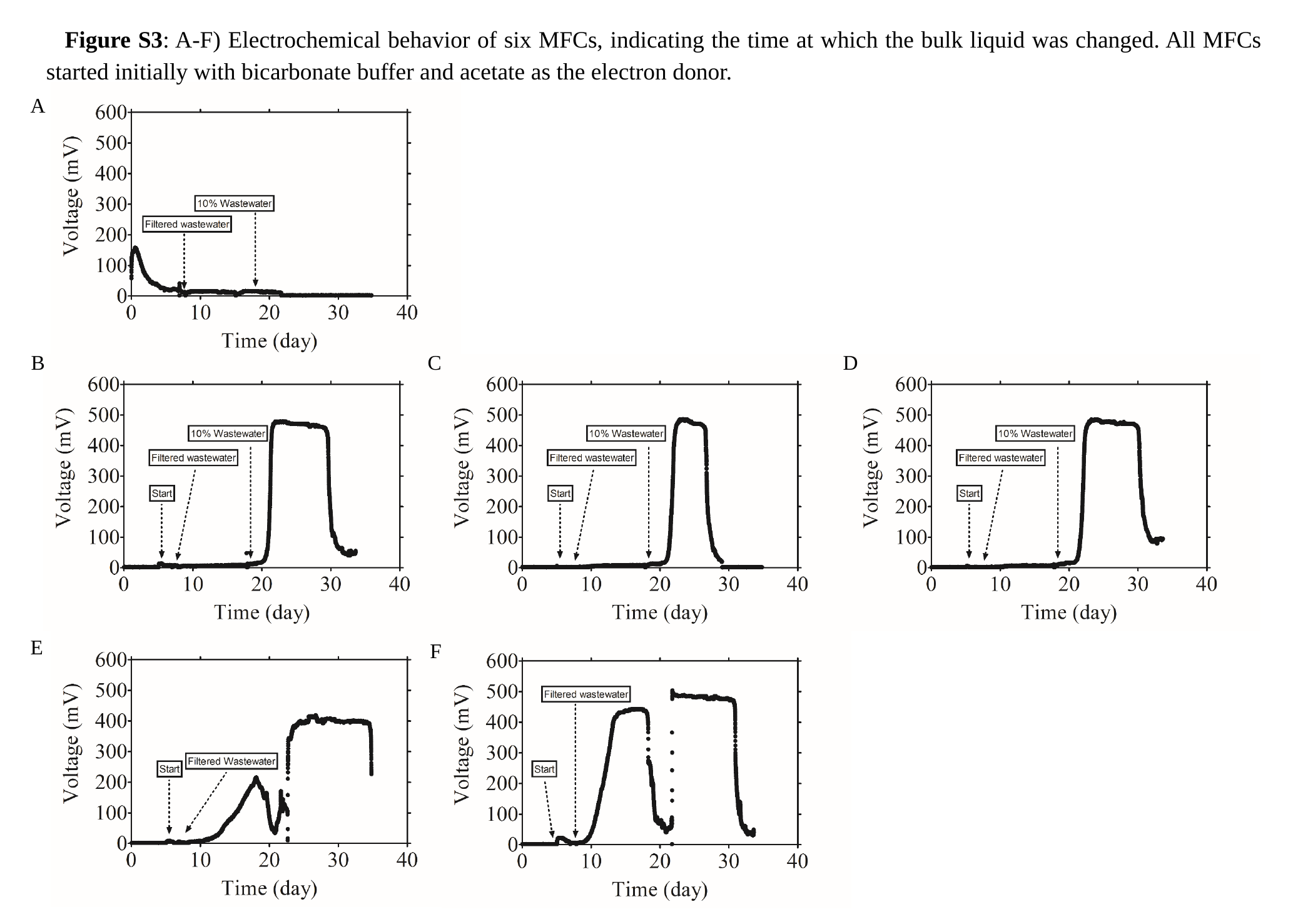

Figure S3: A-F) Electrochemical behavior of six MFCs, indicating the time at which the bulk liquid was changed. All MFCs started initially with bicarbonate buffer and acetate as the electron donor.
A
B
C
D
E
F
