## Supplementary Figure S4 for "Anode surface bioaugmentation enhances deterministic biofilm assembly in microbial fuel cells"

### Slide 1
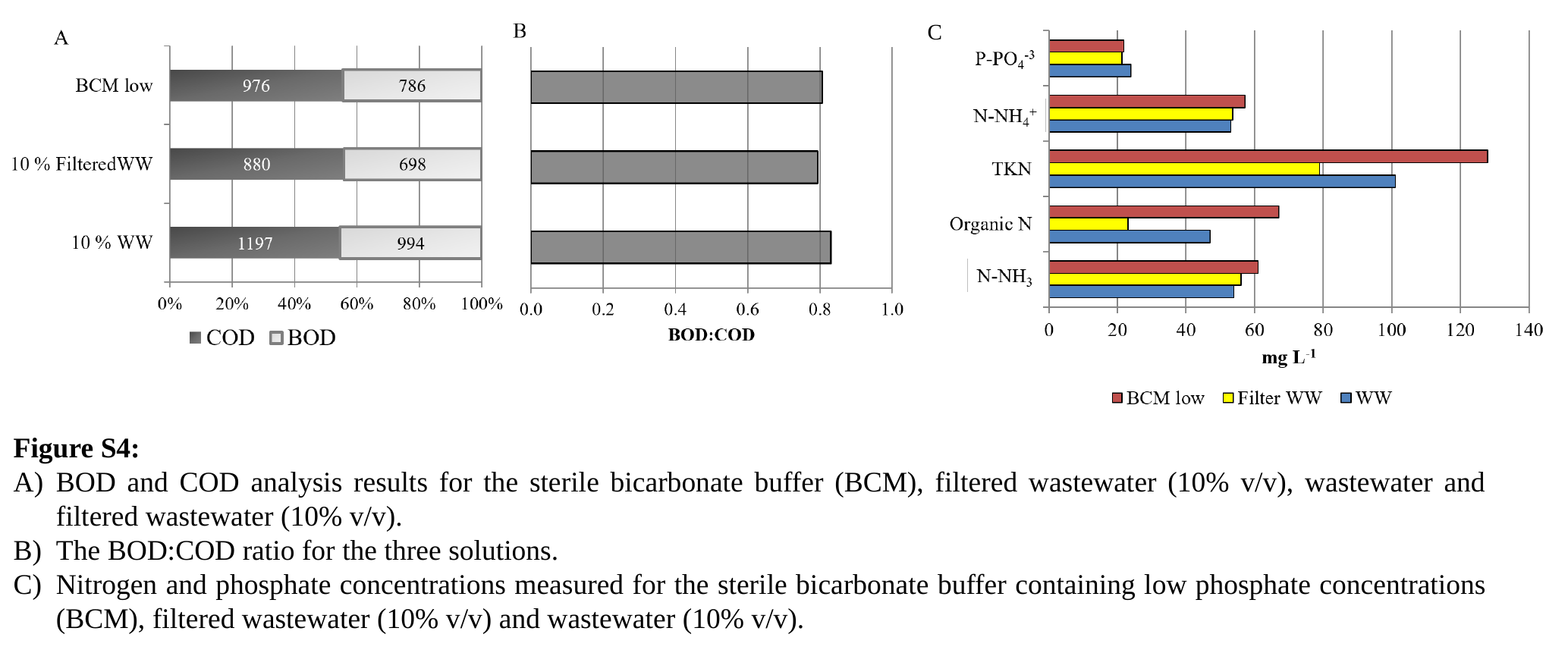

C
Figure S4:
BOD and COD analysis results for the sterile bicarbonate buffer (BCM), filtered wastewater (10% v/v), wastewater and filtered wastewater (10% v/v).
The BOD:COD ratio for the three solutions.
Nitrogen and phosphate concentrations measured for the sterile bicarbonate buffer containing low phosphate concentrations (BCM), filtered wastewater (10% v/v) and wastewater (10% v/v).
