## Supplementary Figure S5 for "Anode surface bioaugmentation enhances deterministic biofilm assembly in microbial fuel cells"

### Slide 1
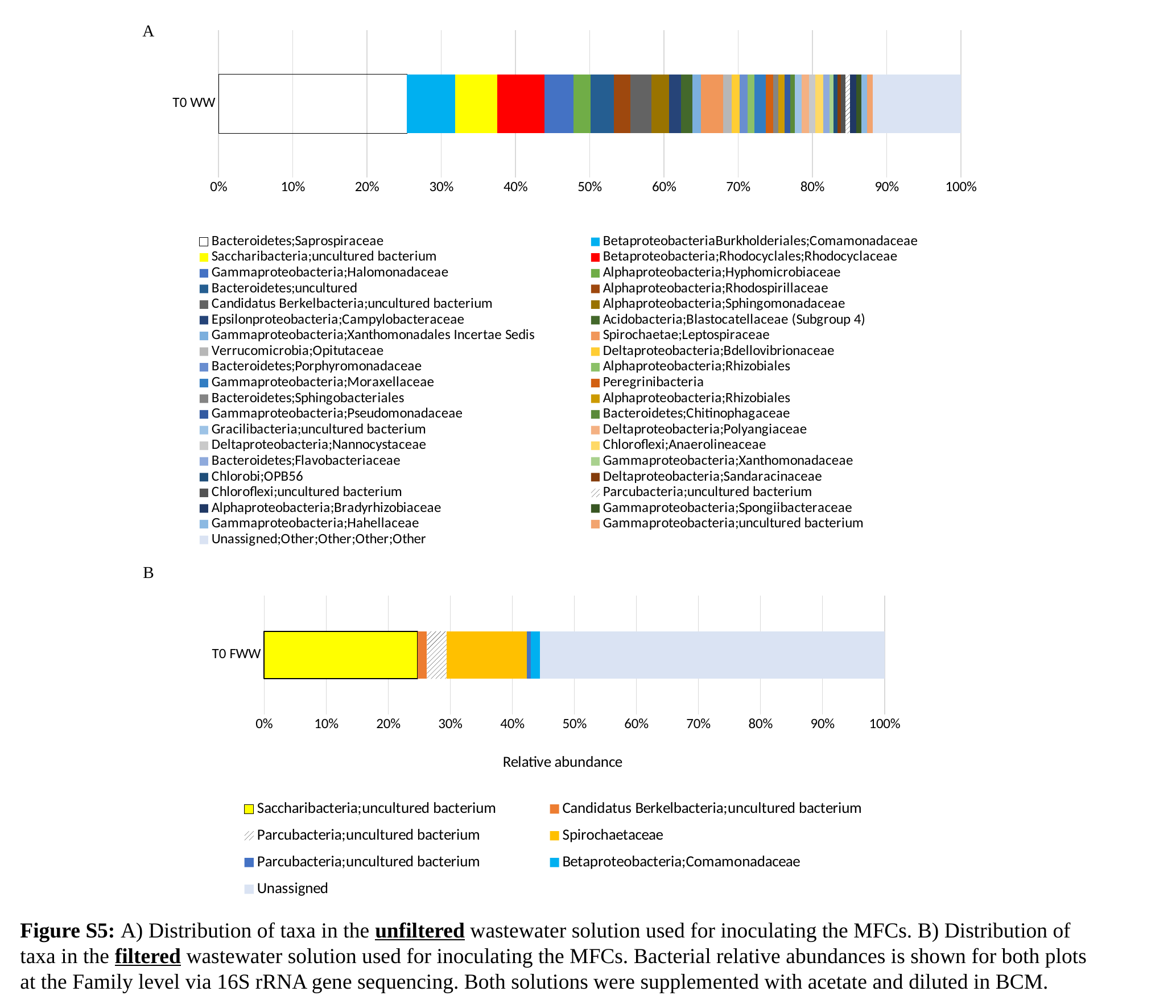

#### Chart
| Category | Bacteroidetes;Saprospiraceae | BetaproteobacteriaBurkholderiales;Comamonadaceae | Saccharibacteria;uncultured bacterium | Betaproteobacteria;Rhodocyclales;Rhodocyclaceae | Gammaproteobacteria;Halomonadaceae | Alphaproteobacteria;Hyphomicrobiaceae | Bacteroidetes;uncultured | Alphaproteobacteria;Rhodospirillaceae | Candidatus Berkelbacteria;uncultured bacterium | Alphaproteobacteria;Sphingomonadaceae | Epsilonproteobacteria;Campylobacteraceae | Acidobacteria;Blastocatellaceae (Subgroup 4) | Gammaproteobacteria;Xanthomonadales Incertae Sedis | Spirochaetae;Leptospiraceae | Verrucomicrobia;Opitutaceae | Deltaproteobacteria;Bdellovibrionaceae | Bacteroidetes;Porphyromonadaceae | Alphaproteobacteria;Rhizobiales | Gammaproteobacteria;Moraxellaceae | Peregrinibacteria | Bacteroidetes;Sphingobacteriales | Alphaproteobacteria;Rhizobiales | Gammaproteobacteria;Pseudomonadaceae | Bacteroidetes;Chitinophagaceae | Gracilibacteria;uncultured bacterium | Deltaproteobacteria;Polyangiaceae | Deltaproteobacteria;Nannocystaceae | Chloroflexi;Anaerolineaceae | Bacteroidetes;Flavobacteriaceae | Gammaproteobacteria;Xanthomonadaceae | Chlorobi;OPB56 | Deltaproteobacteria;Sandaracinaceae | Chloroflexi;uncultured bacterium | Parcubacteria;uncultured bacterium | Alphaproteobacteria;Bradyrhizobiaceae | Gammaproteobacteria;Spongiibacteraceae | Gammaproteobacteria;Hahellaceae | Gammaproteobacteria;uncultured bacterium | Unassigned;Other;Other;Other;Other |
|---|---|---|---|---|---|---|---|---|---|---|---|---|---|---|---|---|---|---|---|---|---|---|---|---|---|---|---|---|---|---|---|---|---|---|---|---|---|---|---|
| T0 WW | 0.20866261400000002 | 0.05334346485714286 | 0.047112462 | 0.052026342428571425 | 0.03206686914285715 | 0.019199594571428568 | 0.025683890428571426 | 0.017983789285714286 | 0.023454913999999997 | 0.019706180428571425 | 0.013221884285714285 | 0.012765957428571428 | 0.009574468 | 0.024468085142857143 | 0.009574468000000001 | 0.008814589714285713 | 0.008763931285714285 | 0.007497466999999999 | 0.012411347428571428 | 0.008156028285714285 | 0.005927051714285715 | 0.006889564285714286 | 0.006484295857142858 | 0.0050658561428571424 | 0.007345491428571428 | 0.008156028428571428 | 0.0071428571428571435 | 0.00871327257142857 | 0.006788247142857142 | 0.0043566361428571425 | 0.004407294714285714 | 0.004154002142857142 | 0.004660587571428571 | 0.005420466285714286 | 0.0069908814285714275 | 0.005673759 | 0.0061296860000000005 | 0.006180344428571427 | 0.09766970614285715 |A
B
#### Chart
| Category | Saccharibacteria;uncultured bacterium | Candidatus Berkelbacteria;uncultured bacterium | Parcubacteria;uncultured bacterium | Spirochaetaceae | Parcubacteria;uncultured bacterium | Betaproteobacteria;Comamonadaceae | Unassigned |
|---|---|---|---|---|---|---|---|
| T0 FWW | 0.24148936166666668 | 0.014420804 | 0.03108747033333333 | 0.12624113466666667 | 0.0055555556666666665 | 0.014775413666666666 | 0.5417257683333333 |Figure S5: A) Distribution of taxa in the unfiltered wastewater solution used for inoculating the MFCs. B) Distribution of taxa in the filtered wastewater solution used for inoculating the MFCs. Bacterial relative abundances is shown for both plots at the Family level via 16S rRNA gene sequencing. Both solutions were supplemented with acetate and diluted in BCM.
