## Supplementary Figure S6 for "Anode surface bioaugmentation enhances deterministic biofilm assembly in microbial fuel cells"

### Slide 1
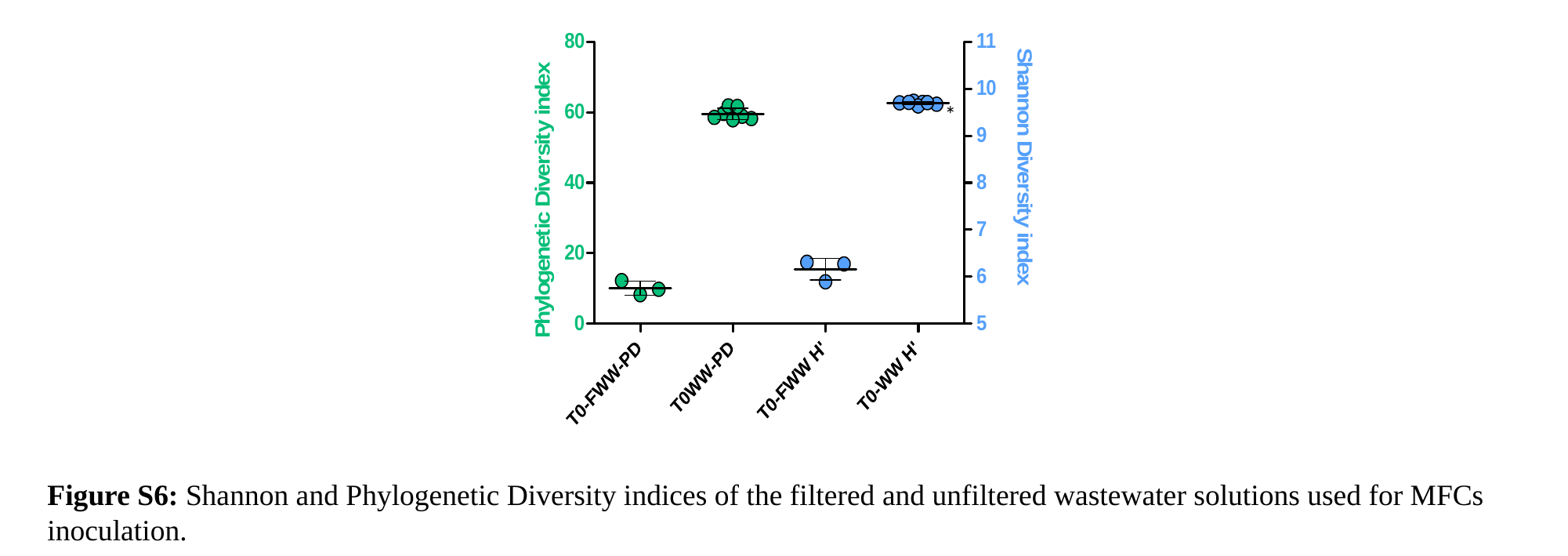

*
*
Figure S6: Shannon and Phylogenetic Diversity indices of the filtered and unfiltered wastewater solutions used for MFCs inoculation.
