## Supplementary Figure S7 for "Anode surface bioaugmentation enhances deterministic biofilm assembly in microbial fuel cells"

### Slide 1
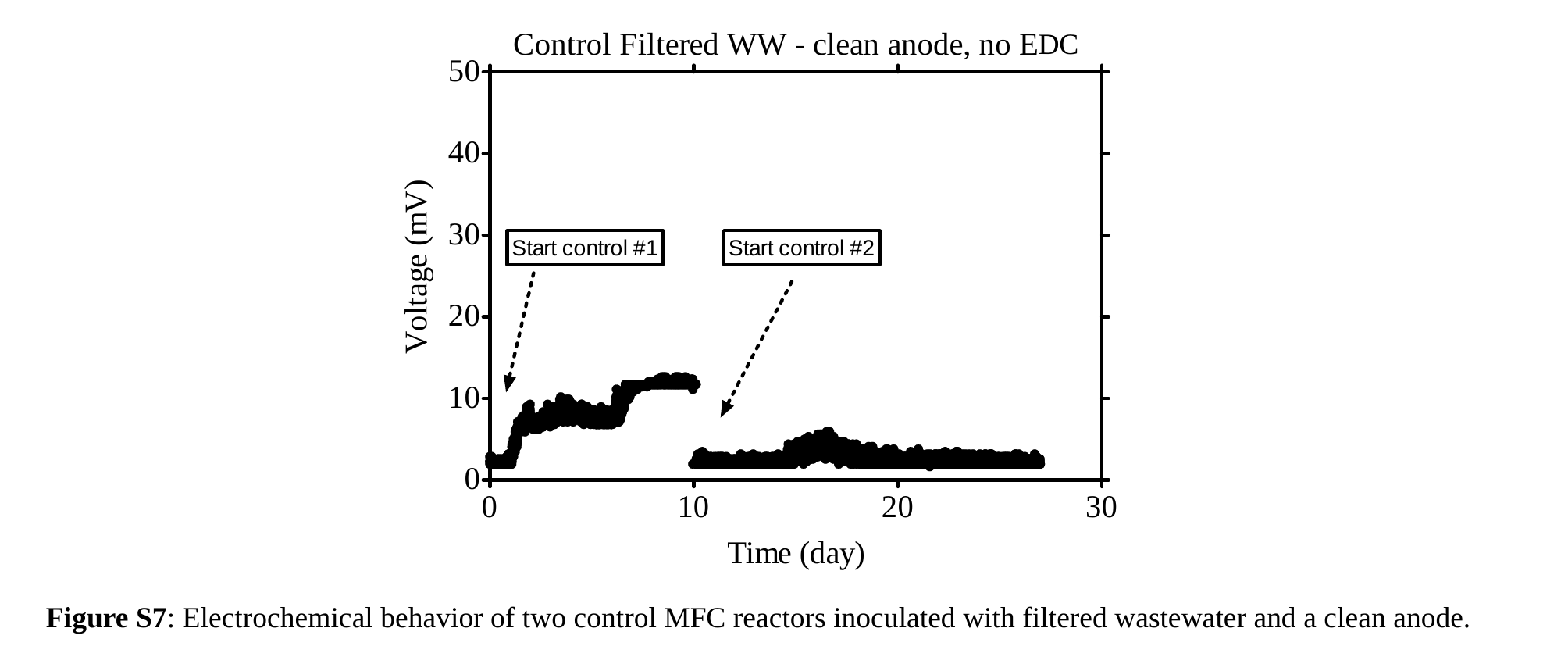

Figure S7: Electrochemical behavior of two control MFC reactors inoculated with filtered wastewater and a clean anode.
DC
