## Supplementary Figure S8 for "Anode surface bioaugmentation enhances deterministic biofilm assembly in microbial fuel cells"

### Slide 1
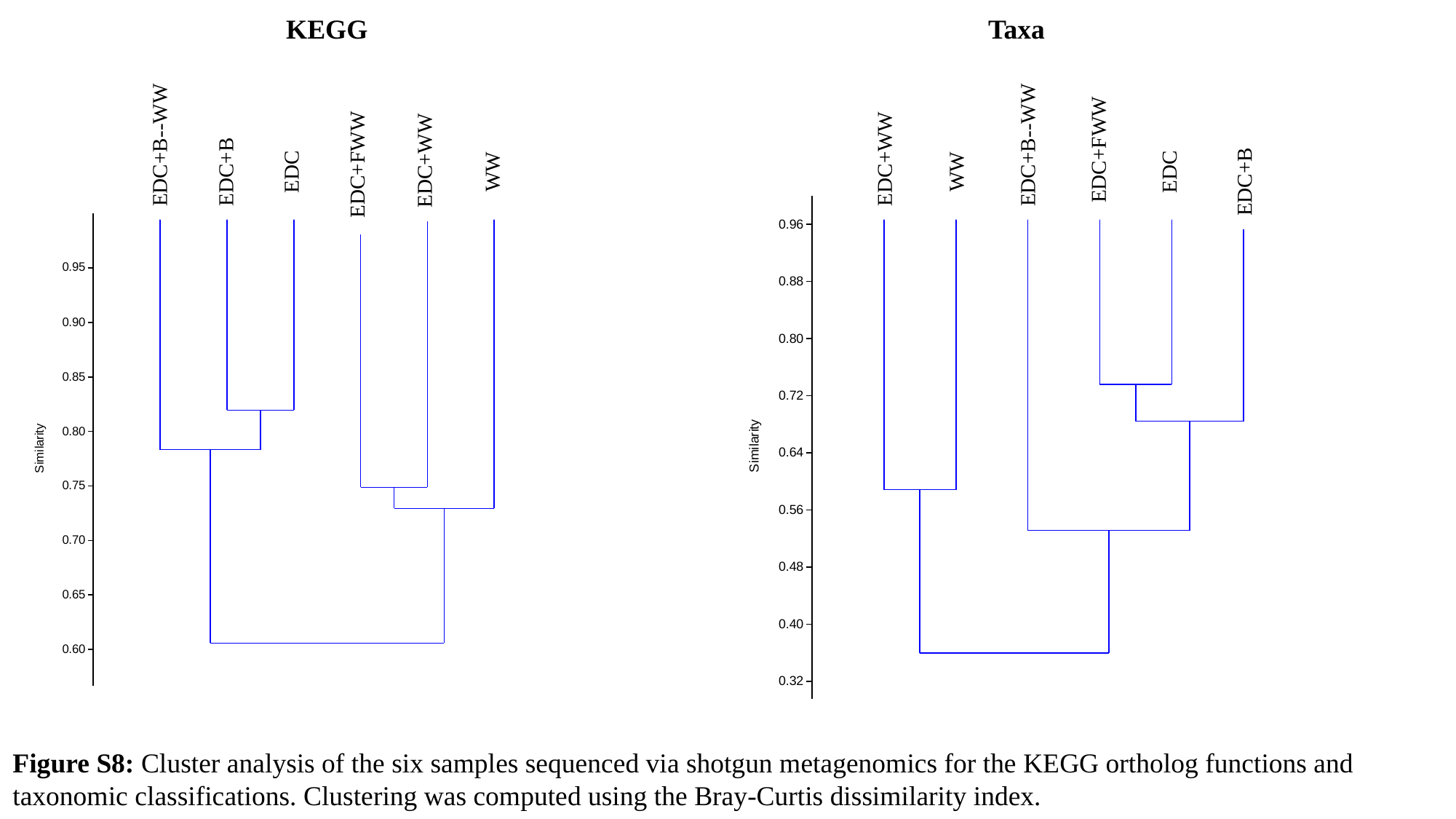

KEGG
Taxa
EDC+B--WW
EDC+WW
EDC+FWW
WW
EDC+B
EDC
EDC+B--WW
EDC+FWW
EDC+WW
WW
EDC
EDC+B
Figure S8: Cluster analysis of the six samples sequenced via shotgun metagenomics for the KEGG ortholog functions and taxonomic classifications. Clustering was computed using the Bray-Curtis dissimilarity index.
