## Supplementary Figure S9 for "Anode surface bioaugmentation enhances deterministic biofilm assembly in microbial fuel cells"

### Slide 1
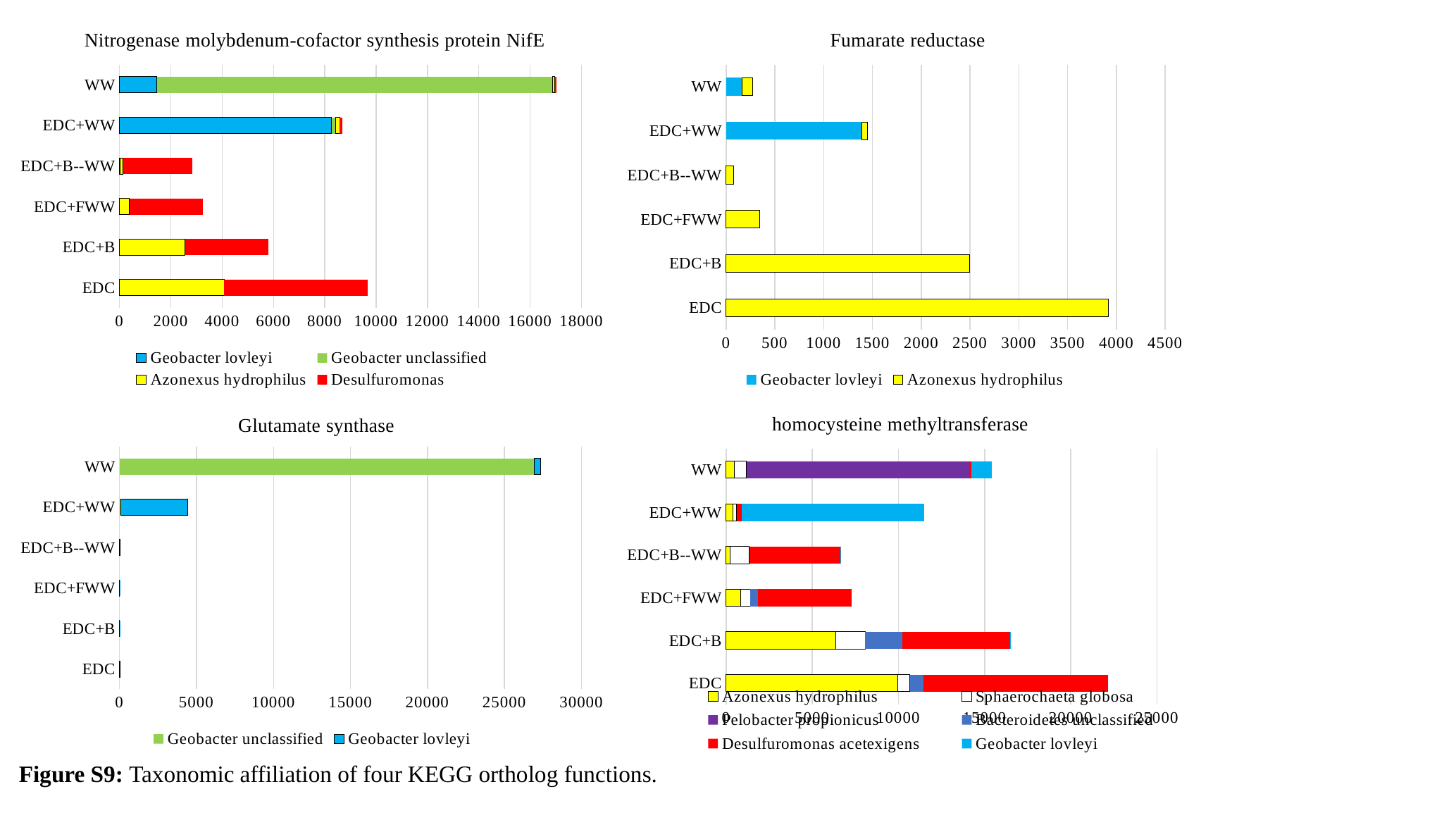

#### Chart: Fumarate reductase
| Category | Geobacter lovleyi | Azonexus hydrophilus |
|---|---|---|
| EDC | 0.0 | 3918.0 |
| EDC+B | 0.0 | 2498.0 |
| EDC+FWW | 0.0 | 346.0 |
| EDC+B--WW | 0.0 | 79.0 |
| EDC+WW | 1389.0 | 58.0 |
| WW | 167.0 | 102.0 |
#### Chart: Nitrogenase molybdenum-cofactor synthesis protein NifE
| Category | Geobacter lovleyi | Geobacter unclassified | Azonexus hydrophilus | Desulfuromonas |
|---|---|---|---|---|
| EDC | 0.0 | 0.0 | 4072.0 | 5599.0 |
| EDC+B | 0.0 | 0.0 | 2556.0 | 3256.0 |
| EDC+FWW | 1.0 | 2.0 | 387.0 | 2872.0 |
| EDC+B--WW | 19.0 | 0.0 | 126.0 | 2705.0 |
| EDC+WW | 8259.0 | 177.0 | 139.0 | 112.0 |
| WW | 1455.0 | 15411.0 | 96.0 | 58.0 |
#### Chart: Glutamate synthase
| Category | Geobacter unclassified | Geobacter lovleyi |
|---|---|---|
| EDC | 11.0 | 6.0 |
| EDC+B | 2.0 | 10.0 |
| EDC+FWW | 7.0 | 4.0 |
| EDC+B--WW | 9.0 | 10.0 |
| EDC+WW | 69.0 | 4375.0 |
| WW | 26920.0 | 420.0 |
#### Chart: homocysteine methyltransferase
| Category | Azonexus hydrophilus | Sphaerochaeta globosa | Pelobacter propionicus | Bacteroidetes unclassified | Desulfuromonas acetexigens | Geobacter lovleyi |
|---|---|---|---|---|---|---|
| EDC | 9966.0 | 698.0 | 6.0 | 785.0 | 10723.0 | 1.0 |
| EDC+B | 6363.0 | 1700.0 | 0.0 | 2162.0 | 6248.0 | 4.0 |
| EDC+FWW | 855.0 | 547.0 | 2.0 | 452.0 | 5426.0 | 0.0 |
| EDC+B--WW | 226.0 | 1115.0 | 0.0 | 4.0 | 5272.0 | 23.0 |
| EDC+WW | 406.0 | 206.0 | 3.0 | 1.0 | 287.0 | 10599.0 |
| WW | 470.0 | 719.0 | 12922.0 | 3.0 | 120.0 | 1198.0 |Figure S9: Taxonomic affiliation of four KEGG ortholog functions.
