## Supplementary Figure S10 for "Anode surface bioaugmentation enhances deterministic biofilm assembly in microbial fuel cells"

### Slide 1
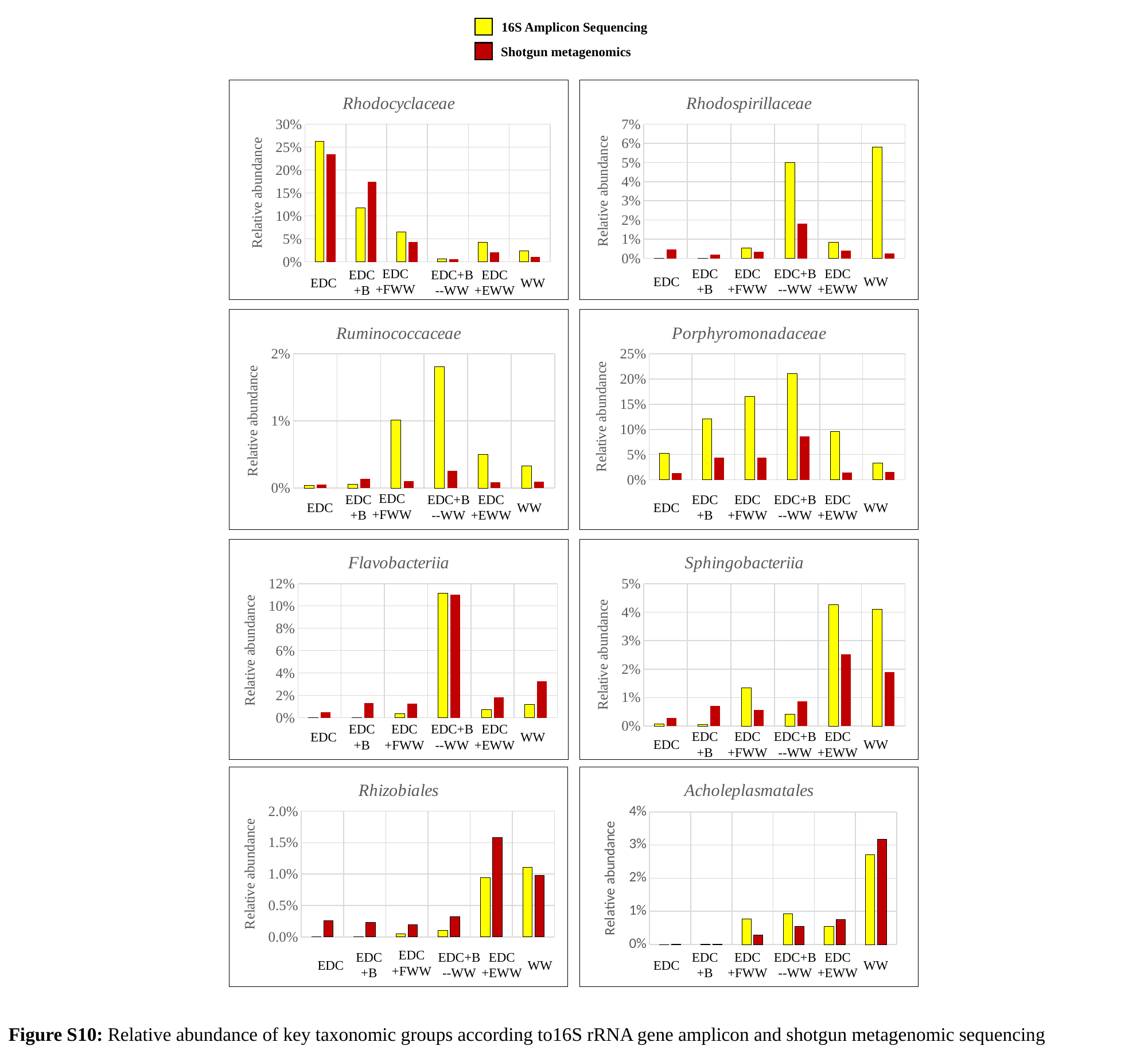

16S Amplicon Sequencing
Shotgun metagenomics
#### Chart: Rhodocyclaceae
| Category | 16S amplicon-seq | SGM |
|---|---|---|
| EDC | 0.263120567375878 | 0.23491314484916742 |
| EDC+B | 0.11713947990544067 | 0.1743518105387582 |
| EDC+FWW | 0.0651773049645484 | 0.043817632484651224 |
| EDC+B--WW | 0.006028368794328 | 0.005553876888947531 |
| EDC+WW | 0.042836879432622 | 0.02083411252391741 |
| WW | 0.023492907801409998 | 0.010844439872653124 |
#### Chart: Rhodospirillaceae
| Category | 16S amplicon-seq | SGM |
|---|---|---|
| EDC | 0.0 | 0.004633986334244998 |
| EDC+B | 0.0 | 0.00188619687579571 |
| EDC+FWW | 0.005390070921977999 | 0.003544333665340231 |
| EDC+B--WW | 0.05 | 0.018004376499610134 |
| EDC+WW | 0.0083687943262464 | 0.0039053706699333536 |
| WW | 0.058067375886655 | 0.0025109536379427366 |EDC
+FWW
EDC
+B
EDC+B
--WW
EDC
+EWW
EDC
WW
EDC
+B
EDC
+FWW
EDC+B
--WW
EDC
+EWW
EDC
WW
#### Chart: Ruminococcaceae
| Category | 16S amplicon-seq | SGM |
|---|---|---|
| EDC | 0.000354609929078 | 0.0005012166139703451 |
| EDC+B | 0.000591016548462 | 0.0013992809704796085 |
| EDC+FWW | 0.0101418439716288 | 0.00104925640875098 |
| EDC+B--WW | 0.01808510638299 | 0.0025741097642298233 |
| EDC+WW | 0.0049645390070891995 | 0.000890719404336254 |
| WW | 0.0032801418439695005 | 0.0009509743285089237 |
#### Chart: Porphyromonadaceae
| Category | 16S amplicon-seq | SGM |
|---|---|---|
| EDC | 0.052836879432604 | 0.01364348167493442 |
| EDC+B | 0.12086288416074735 | 0.04399894761923672 |
| EDC+FWW | 0.165319148936352 | 0.043881820331285515 |
| EDC+B--WW | 0.21063829787263602 | 0.08596076409843546 |
| EDC+WW | 0.095886524822752 | 0.01474020800974998 |
| WW | 0.032978723404264 | 0.01538146919104674 |EDC
+FWW
EDC
+B
EDC+B
--WW
EDC
+EWW
EDC
WW
EDC
+B
EDC
+FWW
EDC+B
--WW
EDC
+EWW
EDC
WW
#### Chart: Flavobacteriia
| Category | 16S amplicon-seq | SGM |
|---|---|---|
| EDC | 0.0 | 0.005006297717400449 |
| EDC+B | 0.0 | 0.01332729713345965 |
| EDC+FWW | 0.0034042553191539994 | 0.012507064302289548 |
| EDC+B--WW | 0.11134751773 | 0.1098763003277487 |
| EDC+WW | 0.0070212765957428005 | 0.018080891505206233 |
| WW | 0.011968085106391501 | 0.03282188183887532 |
#### Chart: Sphingobacteriia
| Category | 16S amplicon-seq | SGM |
|---|---|---|
| EDC | 0.000709219858156 | 0.002781398768464899 |
| EDC+B | 0.0005910165484626666 | 0.007156787797522411 |
| EDC+FWW | 0.0133333333333332 | 0.005772677840019349 |
| EDC+B--WW | 0.004255319148936 | 0.008743944061013743 |
| EDC+WW | 0.0425531914893824 | 0.025203084725797482 |
| WW | 0.041046099290765 | 0.019016765648035294 |EDC
+B
EDC
+FWW
EDC+B
--WW
EDC
+EWW
EDC
WW
EDC
+B
EDC
+FWW
EDC+B
--WW
EDC
+EWW
EDC
WW
#### Chart: Rhizobiales
| Category | 16S amplicon-seq | SGM |
|---|---|---|
| EDC | 0.0 | 0.0025948096138761483 |
| EDC+B | 0.0 | 0.0022702910340896597 |
| EDC+FWW | 0.0004964539007096 | 0.001962442900716721 |
| EDC+B--WW | 0.00106382978723 | 0.003204527882591152 |
| EDC+WW | 0.009432624113472 | 0.015843892681264526 |
| WW | 0.0110815602836875 | 0.00982551472743656 |
#### Chart: Acholeplasmatales
| Category | 16S amplicon-seq | SGM |
|---|---|---|
| EDC | 0.0 | 3.537725036073289e-05 |
| EDC+B | 0.00011820330969266667 | 8.004547079146958e-05 |
| EDC+FWW | 0.0076595744680859985 | 0.0028506751542501183 |
| EDC+B--WW | 0.00921985815603 | 0.005428654021854136 |
| EDC+WW | 0.0055319148936192 | 0.007555625572519548 |
| WW | 0.027127659574475 | 0.03179123024159046 |EDC
+FWW
EDC
+B
EDC+B
--WW
EDC
+EWW
EDC
+B
EDC
+FWW
EDC+B
--WW
EDC
+EWW
EDC
WW
EDC
WW
Figure S10: Relative abundance of key taxonomic groups according to16S rRNA gene amplicon and shotgun metagenomic sequencing
