## Supplementary Methods for "Anode surface bioaugmentation enhances deterministic biofilm assembly in microbial fuel cells"

**Chemical Measurement**

Water samples were filtered prior to the chemical analysis, unless stated otherwise (Amicon Ultra 0.5 ml 10K, Mercury LTD., Israel). Ammonium concentrations were determined with the Sodium Salicylate procedure for total ammonia-nitrogen determination (0-2 mg L^-1^) as modified from APHA (Krom, 1980). Phosphate concentrations were measured with the Vanado Molybdate procedure for high phosphate concentrations (1.0-20 mg L^-1^) followed by standard methods (APHA, 2005). For total COD (Chemical Oxygen Demand) measurements of samples taken at the beginning and at the end of electrochemical cycles, effluent samples were immediately acidified with sulfuric acid (to obtain pH = ~3-4) and stored in 4°C till analyzed. COD was tested based on standard methods (Aminolab Ltd., Israel) or by using the COD HACH kit (HACH COD reactor 45600-00/Hach DR 2010 spectrophotometer, colorimetric method 5220 D). The pH of the bulk liquid was measured by a pH meter (Eutech CON 700, Thermo Scientific) and the conductivity of the bulk liquid was measured by a conductivity meter (Multimeter NM 40, Crison).

**DNA extraction, 16S rDNA amplicon sequencing and shotgun metagenomic sequencing**

Anodic biofilms and planktonic cells from the MFC systems were sampled at different time points for community and phylogenetic analysis, as previously described (Yanuka-Golub et al., 2016). Briefly, total DNA was extracted from anodic biofilms or from the suspended cells using the PowerSoil DNA isolation kit (MO BIO Laboratories) according to the manufacturer’s instructions. For anode sampling, four identical samples were cut out of the carbon felt (0.2 cm depth each disk) and placed directly into the Power Soil® bead tube for further extraction according to the manufacturer’s instructions. For cell suspension, 1300 μl were sampled from the MFC bulk liquid, centrifuged and the pellet was placed directly into the Power Soil® bead tube for further extraction.

For community analysis, a portion of the bacterial 16S rRNA gene was amplified while incorporating linkers required later for sequencing on an Illumina MiSeq sequencer. Bacterial DNA was amplified with the universal 16S rRNA primers targeting the hyper-variable regions V3 and V4, 341F (5'-ACACTGACGACATGGTTCTACANNNNCCTACGGGAGGCAGCAG-3') and 806R (5’-TACGGTAGCAGAGACTTGGTCTGGACTACHVGGGTWTCTAAT-3') (Bartram et al., 2011) under the following conditions: an initial denaturation step at 95°C for 3 min followed by 25 cycles of 95°C for 15 sec, 53°C for 15 sec, and 72°C for 15 sec and ended with an extension step at 72°C for 30 sec. Amplification was verified by electrophoresis on a 1% agarose gel. PCR products were shipped to the DNA Services (DNAS) Facility at the Research Resources Center, University of Illinois at Chicago (UIC), USA, for library preparation and deep amplicon sequencing on an Illumina MiSeq platform. Raw amplicon sequences were merged, quality filtered and trimmed to > 380bp base pairs. Also, the 341F-806R primer sequences were removed during the trimming process by the sequencing service provider. The final FASTA files obtained after quality filtering were further analysed using the QIIME pipeline, version 1.8 (Caporaso et al., 2010) and chimeric sequences were removed with the UCHIME algorithm implemented within USEARCH (Edgar et al., 2011). Sequences were clustered to OTUs (operational taxonomic unit) at 97% similarity (or 99% when was necessary) using the UPARSE algorithm (Edgar, 2013), then, representative sequences and an OTU table were generated accordingly. Sequences were assigned taxonomically using the UCLUST algorithm (Edgar, 2010) against the SILVA 16S rRNA database (Quast et al., 2012). The number of sequences per sample was rarefied to an equal depth to avoid a biodiversity bias due to unequal sampling effort.

Genomic libraries were prepared with the Nextera XT library preparation kit using approximately 1 ng total DNA per sample (DNA concentrations verified by Qubit fluorometry). Shotgun metagenomic sequencing was done using Illumina NextSeq500 paired-end (2 x 150 bp reads) at DNA Services Facility, as well. Six anode samples were successfully sequenced and processed for all the downstream analysis. The sequencing depth (after quality control step) for the samples was as follows: **EDC** = 5.2 Gbp, **EDC+B** = 4.9 Gbp, **EDC+B**--WW = 5.8 Gbp, **EDC+**WW = 5.5 Gbp, **WW** = 10.4 Gbp and **EDC+**FWW = 2.7 Gbp. Each of the five anode samples was taken after the MFC reactor completed one electrochemical cycle (Supplementary Figure S3G). Quality control step was done with Trimmomatic v0.36 [N1] using default parameters, which consisted of low-quality reads filtering and removal of low score bases and short reads.

**Metagenomic assembly analysis**

*De novo* metagenome assemblies were done using SPAdes assembler v3.11.0 [N2]; In order to improve assembly by increasing sequenced reads coverage, samples **EDC+B**, **EDC+B**--WW and **EDC+**FWW were co-assembled, **EDC+WW** and **WW** were co-assembled and sample **EDC** was assembled alone. The rationale behind this specific samples separation into three assemblies was the bacterial diversity and compositional similarity between the samples (analyzed prior with 16S rRNA sequencing). SPAdes was run in paired-end mode with the following parameters: --careful (to minimize the number of mismatches in the contigs), -k 21, 33, 55, 77, 99, 127 (k-mer lengths). A summary of the metagenome assemblies’ statistics is found in Supplementary Table 1. Assembled contigs smaller than 1kb were discarded. The rest of the contigs were used for open reading frames prediction using the gene finding algorithm Prodigal v2.6.3 [N3]. In order to create one unified dataset of genes with reduced redundancy, the predicted proteins from the three assemblies were combined and clustered at 90% identity using CD-HIT v4.6 [N4]. To quantify the predicted genes for each sample, the metagenomic reads of each sample were separately mapped against the unified and clustered genes set using Bowtie2 short reads mapper v2.2.9 [N5] in --very-sensitive-local mapping mode and SAMtools v1.3.3 [N6]. Next, to filter out low coverage genes, a threshold cutoff of 100 reads summed across all six samples was applied. After the 90% protein clustering and 100 reads cutoff, the size of the unified protein set was reduced by 75% (from 792,893 to 192,446) but retained 92.7±7.9% (mean±sd) of the total reads in each sample (for all clustering and threshold cutoff mapping results see Supplementary Table 2).

Finally, to obtain functional and taxonomic annotations for the proteins set, BLASTP search against NCBI reference proteins (RefSeq) database (downloaded May 15^th^, 2017) was done using DIAMOND [N7], run with default parameters except --max-target-seqs set to 50 (up to 50 hits per read). For taxonomic assignments of the reads, files were imported into MEGAN5 (<http://ab.inf.uni-tuebingen.de/software/megan5/>), and functional annotation of the reads was done based on the KEGG library (Kyoto Encyclopedia for Genes and Genomes, <http://www.genome.jp/kegg/>).

To assemble the genome of *Desulfuromonas* from the metagenomic samples, the genome sequence of *D. acetexigens* (the species most closely related to the *Desulfuromonas* enriched in the *Desulfuromonas* enriched consortium) from Katuri et al., 2017 [N8] was used as a reference genome. More specifically, reads from EDC+B, EDC+B—WW, EDC+FWW and EDC samples were separately mapped to the reference genome (EDC+WW and WW samples had too low number of reads mapped to the reference genome to obtain a decent assembly) using Bowtie2 as described above in order to separate these reads from the rest of the metagenome. Next, the mapped reads were used as an input for genome assembly with SPAdes as described above. Assembled contigs smaller than 500bp were discarded. The assembled genomes completeness, contamination and strain heterogeneity levels were analyzed with CheckM v1.0.7 software [N9] in specific taxon marker set mode (taxonomy_wf) and are reported in Supplementary Table 3 (CheckM_output.xlsx). Average nucleotide identity (ANI) between each genome pair from this study and with the reference genome was calculated with OrthoANI software (OAU v1.2) [N10].	Metagenomic sequence data generated in this study have been deposited to NCBI SRA under BioProject ID PRJNA594175

**Electroactive bacteria enrichment**

A single anode sample (one 0.2 cm depth disk) was placed into a 2 ml Eppendorf filled with BCM medium and 4-5 sterilized glass beads. Sample was vortexed at half of maximal speed, on the flat-bed adaptor, for 2.5 minutes. At this point anode "dust" was observed but most of the anode remained whole. 100μl of each crude sample and of a x10 dilution (in BCM) were plated on BCM-Na_2_S and BCM-Sodium Fumarate plates (Supp. Table 4). Plates were kept in an anaerobic jar at room temperature in the dark. Orange-colored colonies appeared after 3 weeks and were re-plated on the BCM-Sodium Fumarate media plate. For taxonomic identification, the full 16S rDNA gene of selected colonies was amplified using universal 16s rRNA primers (primer set 8F-1512R), sequenced and compared to NCBIs nr database using blastn. For each MFC inoculation, in order to directly apply the isolate on the anode, all the consortium growing on a single plate was harvested using an inoculation loop and spread over one side of the anode. The *Desulfuromonas* enriched consortium was continuously re-plated to allow several MFC inoculation attempts and replicates.

Finally, to assure a consistent appearance of the same *Desulfuromonas* species that was applied to the MFCs after enrichment, the *D. acetexigens* genome’s sequence (Katuri et al., 2017) was used as a reference to assemble this genome separately from each of the anode samples collected once steady-state current production was achieved: EDC, EDC+B, EDC+B--WW and EDC+FWW (in samples WW and EDC+WW the number of reads that matched *D. acetexigens* was too low to assemble a draft genome). The completeness levels of the assembled genomes ranged from 83-91%, however, contamination (12-19%) and strain heterogeneity (80-90%) were high even after additional analyses (see above), indicating the existence of several closely related strains of *D. acetexigens* within these genomes.
